# ART duration and immunometabolic state determine efficacy of DC-based treatment restoring functional HIV- specific CD8+ T cells in PLWH

**DOI:** 10.1101/2022.01.14.476403

**Authors:** Marta Calvet-Mirabent, Ildefonso Sánchez-Cerrillo, Noa Martín-Cófreces, Hortensia de la Fuente, Ilya Tsukalov, Cristina Delgado-Arévalo, María José Calzada, Ignacio de los Santos, Jesús Sanz, Lucio García-Fraile, Francisco Sánchez-Madrid, Arantzazu Alfranca, María Ángeles Muñoz-Fernández, Maria J. Buzón, Enrique Martín-Gayo

**Affiliations:** Immunology Unit from Hospital Universitario de La Princesa and Instituto de Investigación Sanitaria Princesa, Madrid, Spain; Universidad Autónoma de Madrid, Madrid, Spain; Centro de Investigación Biomédica en Red Cardiovascular, CIBERCV, 28029 Madrid, Spain; Infectious Diseases Unit from Hospital Universitario de La Princesa, Madrid, Spain; Immunology Section, Instituto de Investigación Sanitaria Gregorio Marañón (IiSGM), Hospital General Universitario Gregorio Marañón, Madrid, Spain; Infectious Diseases Department, Institut de Recerca Hospital Univesritari Vall d’Hebrón (VHIR), Universitat Autònoma de Barcelona, Barcelona, Spain; Centro de Investigación Biomédica en Red Infecciosas, CIBERINF, 28029 Madrid, Spain

## Abstract

Dysfunction of CD8+ T cells in people living with HIV-1 (PLWH) receiving anti-retroviral therapy (ART) has restricted the efficacy of dendritic cell (DC)-based immunotherapies against HIV-1. Heterogeneous immune exhaustion and metabolic states of CD8+ T cells might differentially associate with dysfunction. However, specific parameters associated to functional restoration of CD8+ T cells after DC treatment have not been investigated in detail. Here, we studied the association of ART duration with memory subsets, exhaustion and metabolic profiles of CD8+ T cells from PLWH and improvement of polyfunctional and effector HIV-1 specific responses after stimulation with Gag-adjuvant-primed DC. HIV-1-specific CD8+ T cell responses from a larger proportion PLWH on ART for more than 10 years (LT-ARTp) improved polyfunctionality and capacity to eliminate autologous p24+ infected CD4+ T cells *in vitro*. In contrast, CD8+ T cells from PLWH on ART for less than a decade (ST-ARTp) were less responsive to DC treatment and functional improvement was limited in this group. This was associated with lower frequencies of central memory CD8+ T cells, increased co-expression of PD1 and TIGIT and reduced mitochondrial respiration and glycolytic induction upon TCR activation. In contrast, CD8+ T cells from LT-ARTp showed increased frequencies of TIM3+PD1-cells and preserved induction of glycolysis. Treatment of dysfunctional CD8+ T cells from ST-ARTp with combined anti-PD1 and anti-TIGIT antibodies plus a glycolysis promoting drug restored their ability to eliminate infected CD4+ T cells. Together, our study identifies specific immunometabolic parameters for different PLWH subgroups potentially useful for future personalized DC-based HIV-1 vaccines.

## INTRODUCTION

Current available antiretroviral therapies (ART) do not eradicate chronic HIV-1 infection due to persistent HIV-1 reservoir maintained through homeostatic proliferation of long-lived latently infected memory CD4+ T cells, the residual viral replication in tissues and dysfunctionality of the immune response (1-6). Under these conditions, it would take over 70 years to eliminate the HIV reservoir with the actual ART (7). Therefore, novel strategies aiming to eradicate the latent reservoir continue to be investigated. During the last decade, the field has focused on *shock and kill* strategies, based on the viral reactivation of latently HIV-1 infected cells using latency reversal agents (*shock*) and their elimination by improved immune cytotoxic cell subsets (*killing*) (8). In this regard, cytotoxic CD8+ T lymphocytes (CTL) are crucial immune cells for restraining the size of the reservoir in the initial phases of the infection (9) and for natural control of HIV-1 replication in elite controller (EC) individuals (10). Several therapeutic strategies have focused on enhancing the magnitude and polyfunctionality of HIV-specific CD8+ T cell responses of people living with HIV-1 (PLWH) (10-12) to mimic effective and durable CTL responses of EC (12). However, none of the tested candidates for CD8+ T cell prevented viral rebound following treatment interruption in clinical trials (13, 14). These negative results suggest the need of more personalized strategies which take into consideration complex factors contributing to CD8+ T cell dysfunction in these individuals.

A potential strategy is to improve antigen-presenting cell function of dendritic cells (DC), which could also be compromised in PLWH (15). In this regard, previous studies on HIV-1 infected EC identified a DC activation state dependent on the TANK-Binding Kinase 1 (TBK-1) characterized by highly functional capacities to activate polyfunctional HIV-specific CD8+ T cell responses (16, 17). TBK-1 is a master regulator downstream of multiple intracellular nucleic acid sensors such as TLR3/TRIF, RIG-I/MDA5 and cGAS/STING, which leads to the production of type I and II interferons (18). Therefore, activation of DC through agonists of nucleic acid sensors upstream of TBK-1 could be useful to potentiate their function to an EC-like phenotype. Consistently, our previous studies suggested that vaccination of humanized mice with DC primed with TBK1-adyuvants increased polyfunctional HIV-1-specific CTL responses in lymphoid tissues (19). In addition, the use of DC might also be intrinsically useful to more efficiently reactivate the latent HIV-1 reservoir (20, 21).

On the other hand, basal hyperactivation and immune exhaustion compromise the maintenance, functionality and effector function of CD8+ T cells against HIV-1-infected cells in PLWH (22-25). Multiple recent studies have shown that exhaustion of CD8+ T cells is a dynamic process involving expression of multiple checkpoint inhibitory receptors that drive the transition, survival and function of distinct memory cell subsets (24, 26, 27). Importantly, expression of checkpoint inhibitory receptors such as PD1, TIGIT and TIM3 in T cells is also associated with deregulated glucose metabolism (28-30), which could lead to impaired effector function and longevity of HIV-1 specific CD8+ T cells (31-33). In fact, compromise of metabolic fitness has been reported in PLWH and suggested to affect CD8+ T cell function (34, 35). However, it is unknown whether specific patterns of expression of checkpoint inhibitory receptors might differentially associate with efficacy of DC treatment inducing polyfunctionality and cytotoxic ability of HIV-1-specific CD8+ T cells in PLWH on ART.

In the present study, we evaluated the efficacy of adjuvant-primed DC reinvigorating polyfunctional and cytotoxic competent HIV-1-specific CD8+ T cells, and the association of this response with the immunometabolic state in different subgroups of PLWH at different times since ART initiation. We report enhanced response of CD8+ T cells to DC treatment in PLWH on ART for more than a decade, which associates with higher central memory (CM) frequencies, enrichment on TIM3+ PD1-cells and the ability to induce glycolysis upon TCR stimulation. In contrast, CD8+ T cells from PLWH under ART for less than 10 years were characterized by decreased frequency of CM, and an increased proportions of exhausted PD1+ TIGIT+ memory CD8+ T cells. These patterns correlated with impaired glycolytic metabolism and inability to reinvigorate cytolytic function of CD8+ T cells from these individuals after DC stimulation. Importantly, combined blockade of PD1 and TIGIT receptors and the use of the pro-glycolytic drug metformin more consistently restored functional cytotoxic ability of dysfunctional CD8+ T cells from PLWH. Collectively, our study uncovers new immunometabolic parameters associated with response to DC-based HIV-1 vaccine in PLWH, which might be useful for future personalized treatments.

## MATERIALS AND METHODS

### Study participants and Ethic statement

Human Peripheral blood mononuclear cells (PBMC) were isolated by Ficoll (Pancoll, PAN Biotech) gradient centrifugation from peripheral blood samples of n=49 PLWH (age 48 [25-74], 86% males) on antiretroviral therapy (8 [1-24] years) characterized by undetectable plasma viremia, 866 [430-1676] median CD4+ T cell counts and with clinical characteristics summarized in Table 1 provided by the Infectious Diseases Unit from Hospital de La Princesa, Madrid, Spain. N=12 HIV negative donors (Age 45.5 [23-70], 64% males) were provided either by the Immunology Service from Hospital de La Princesa or Centro de Transfusión of Comunidad de Madrid, Spain (Table 2) and used for comparison purposes. All participating subjects accepted donating samples after receiving and signing an informed consent, approved by the Ethics committee from Hospital Universitario de La Princesa (Register Number 3518) and following the Helsinki declaration.

**Table 1.** Clinical characteristics of ART-treated HIV-1 chronic patient cohort.

|  | Total HIV chronic patient cohort | ST-ARTp | LT-ARTp | p-values |
| --- | --- | --- | --- | --- |
| **Patient cohort individuals (n)** | 49 | 31 | 18 |  |
| **Age at sample collection**<br>Median [min, max];<br>stdev p | 48 [25, 74]; 12 | 40 [25, 57]; 9 | 55.5 [40, 74]; 10 | <0.0001 |
| **Sex**<br>M: male (%);<br>F: female (%) | M: 42 (86%);<br>F: 7 (14%) | M: 30 (97%);<br>F: 1(3%) | M: 12 (67%);<br>F: 6 (33%) | 0.013 |
| **Circulating CD4+ T cell counts at sample collection**<br>Median [min, max];<br>stdev p | 866 [430, 1676]; 330<br>1 ND | 1016 [430, 1676];<br>357<br>1 ND | 777 [544, 1259];<br>225 | 0.039 |
| **Circulating CD8+ T cell counts at sample collection**<br>Median [min, max];<br>stdev p | 900 [416, 2719]; 489<br>1 ND | 993 [493, 2719];<br>527<br>1ND | 833 [416, 1614];<br>383 | ns |
| **CD4+/CD8+ T cell ratio in the blood at sample collection**<br>Median [min, max];<br>stdev p | 1.00 [0.27, 2.44];<br>0.46<br>1 ND | 1.06 [0.27, 1.82];<br>0.41<br>1 ND | 0.97 [0.4, 2.44];<br>0.53 | ns |
| **Plasma viral load** | Undetectable<br>(<20 copies/ml) | Undetectable<br>(<20 copies/ml) | Undetectable<br>(<20 copies/ml) |  |
| **BLIPS**<br>0: patients (%);<br>1: patients (%);<br>ND: patients (%) | 0: 34 (69.4%);<br>1: 8 (16.3%);<br>ND: 7 (14.3%) | 0: 23 (74%);<br>1: 5 (16%);<br>ND: 3 (10%) | 0: 11 (61%);<br>1: 3 (17%);<br>ND: 4 (22%) | ns |
| **HCV co-infection** | No | No | No |  |
| **Years under treatment**<br>Median [min, max];<br>stdev p | 8 [1, 24]; 6 | 5.5 [1, 9.5]; 3 | 15 [10.5, 24]; 3 | <0.0001 |
| **Treatment** [number of patients] | ABC/3TC/DTG [9];<br>ABC/3TC/LPV/r [1];<br>DRV/c [6];<br>DRV/c/DTG/3TC [1];<br>DRV/c/TDF/AZT [1];<br>DRV/RPV/FTC [1];<br>DTG/RPV [1];<br>DTG/3TC [2];<br>TAF/FTC/BIC [5];<br>TAF/FTC/DRV/c [9];<br>TAF/FTC/DRV/c/DTG [1]; TAF/FTC/EVG/c | ABC/3TC/DTG [6];<br>ABC/3TC/LPV/r [0];<br>DRV/c [2];<br>DRV/c/DTG/3TC [0];<br>DRV/c/TDF/AZT [1];<br>DRV/RPV/FTC [0];<br>DTG/RPV [1];<br>DTG/3TC [2]; | ABC/3TC/DTG [3];<br>ABC/3TC/LPV/r [1];<br>DRV/c [4];<br>DRV/c/DTG/3TC [1];<br>DRV/c/TDF/AZT [0];<br>DRV/RPV/FTC [1];<br>DTG/RPV [0];<br>DTG/3TC [0]; |  |
|  | [5]; TAF/FTC/RPV [2];<br>TAF/FTC/NVP [1];<br>TDF/FTC/EFV [4] | TAF/FTC/BIC [5];<br>TAF/FTC/DRV/c<br>[6];<br>TAF/FTC/DRV/c/D<br>TG [0];<br>TAF/FTC/EVG/c<br>[3]; TAF/FTC/RPV<br>[2]; TAF/FTC/NVP<br>[0];<br>TDF/FTC/EFV [3] | TAF/FTC/BIC [0];<br>TAF/FTC/DRV/c<br>[3];<br>TAF/FTC/DRV/c/D<br>TG [1];<br>TAF/FTC/EVG/c<br>[2]; TAF/FTC/RPV<br>[0]; TAF/FTC/NVP<br>[1];<br>TDF/FTC/EFV [1] |  |

\*ND = non determined
Statistical significances were calculated using two-tailed Wilcoxon test and Chi-square test with Yates correction.

**Table 2.** HIV negative donor cohort.

|  |  |
| --- | --- |
| **Healthy control cohort individuals (n)** | 12 |
| **Age at sample collection**<br>Median [min, max]; stdev p | 45.5 [23, 70]; 15 |
| **Sex**<br>M: male (%); F: female (%) | M: 7 (64%);<br>F: 4 (36%) |

### Monocyte-derived dendritic cell (MDDC) generation and in vitro stimulation

Monocyte-derived DCs (MDDCs) were generated from circulating monocytes enriched by adherence and cultured in RPMI 1640 media supplemented with 10 % Fetal Bovine Serum (HyClone) in the presence of 100 IU/ml of GM-CSF and 100 IU/ml of IL4 (Prepotech) for 6 days. Subsequently, MDDCs were harvested and cultured over-night in media alone or in the presence of 5 μg/ml of a pool of 15-mer overlapping HIV-1 Gag peptides provided by the NIH AIDS Reagent Program (#11057) either individually or in combination with 1 μg/ml of 2’3’-c’diAM(PS)2 (Invivogen) STING agonist and 2.5 μg/ml of Poly I:C (SIGMA) TLR3 ligand as adjuvant treatment. Maturation of MDDC after treatment with Gag-peptides and/or adjuvants was assessed by flow cytometry analysis of CD40 expression in these cells.

### Isolation of human peripheral blood T lymphocyte populations

Total or individual CD8+ or CD4+ T cells were purified from PBMC suspensions from our PLWH and HIV-1 negative control study cohorts by negative immunomagnetic selection (purity >90%) using the Untouched Human T cell and CD4 T cell Dynabeads Kits (Invitrogen) and the MojoSort Human CD8 T cell (BioLegend) isolation kit. In some experiments, memory CD45RA-CD8+ T cells from PLWH and HIV-1 negative controls were also isolated by sequential negative immunomagnetic selection protocol. Briefly, total CD8+ T cells were isolated from PBMCs as previously described and subsequently, memory cells were purified after treatment with mouse anti-human CD45RA mAbs (Biolegend) and Goat anti-Mouse IgG covered Dynabeads (Invitrogen).

### In vitro activation of HIV-1 specific CD8+ T cell responses from PLWH

To assess the impact of MDDC therapy on the magnitude and polyfunctionality of HIV-1 specific CD8+ T cell responses, MDDCs pretreated with media only or either with the Gag peptides alone or incubated with adjuvants were co-cultured with purified autologous T cells in a 1:5 ratio. After 5 h incubation, 0.25 μg/ml Brefeldin A (SIGMA), 0.0025 mM Monensin (SIGMA) and 0.2 μg/ml anti-CD107a-APC antibody (Biolegend) were added to the media and then cultured over-night. Analysis of magnitude (total IFNγ expression) and polyfunctionality (co-expression of IFNγ and CD107a) of CD8+ T cells after MDDC and antigen stimulation was analyzed by flow cytometry.

### In vitro analysis of CD8+ T cells functionality

To evaluate cytotoxic function of CD8+ T cells from PLWH on ART receiving DC therapy, CD8+ T cells were precultured with autologous MDDC as previously described at a 1:3 ratio. After 24 h, unstimulated CD4+ T cells were purified from the same patient, stained with 5 μM violet cell trace tracker (Invitrogen), and cultured in the presence of 30 μM Raltegravir (Selleck Chemicals) alone or in combination with 50 nM Romidepsin (Selleck Chemicals) in the absence or the presence of autologous CD8+ T cells treated with MDDC pretreated under different conditions at a 2:1 ratio. After 24 h, intracellular HIV-1 p24 protein levels present in violet+ CD4+ T cells were analyzed by flow cytometry. In some functional experiments, we evaluated the impact of blocking different check point inhibitory receptors and/or stimulating glycolysis in CD8+ T cells during co-culture with the activated MDDCs. In these assays, media was supplemented with 2 μg/ml of mouse IgG1 K anti-human PD-1 antibody (Biolegend) either alone or in combination with 1 μg/ml mouse IgG2 B anti-human TIGIT (R&D systems) or 1 μg/ml goat IgG anti-human TIM-3 (R&D systems) and alone or in combination of 5 mM Metformin (SIGMA). As a control, we used the appropriate corresponding isotype control antibodies (mouse IgG1 K and mouse IgG2 B, both from Biolegend, goat IgG from SIGMA) and a drug carrier control.

### Flow cytometry

Analysis of cell viability *ex vivo* or after culture was performed using APC-H7 (Tonbo Biosciences) or LIVE/DEA Fixable Yellow 405 (invitrogen) viability dye, in the presence of different panels of monoclonal antibodies. For analysis of MDDC activation we used Lineage (CD3 APC, CD19 APC, CD56 APC), CD11c Pacific Blue, CD40 FITC, and HLA-DR PerCP (all from Biolegend). For analysis of *in vitro* activation of HIV-1 specific CD8+ T cell responses anti-human CD3 CF-Blue (Immunostep), CD8 PerCP, CD107a APC (Biolegend) and IFNγ FITC (BD Pharmigen) mAbs were used. For functional assays with CD8+ T cells anti-human CD3 FITC, CD4 APC, CD8 PerCP (Biolegend), and anti-HIV-1 p24 PE (Beckman coulter) were used. CD8+ T cell memory and exhaustion phenotypes were defined using anti-human CD3 CF-Blue (Immunostep), CD8 APC/Cy7, CD45RO PE/Cy7, PD1 PerCP, TIGIT APC and TIM3 FITC (Biolegend), and CCR7 PE (R&D systems). Clones of antibodies used in our study are detailed in Table 3. For intracellular staining we used Fixation buffer and Intracellular staining Perm Wash buffer (Biolegend). Samples were analyzed on a FACS Canto II cytometer (BD Biosciences). Analysis of individual and Boolean multiparametric flow cytometry data was performed using FlowJo v10.6 software (Tree Star).

**Table 3.** List of commercial antibodies used in the study

| Antibody | Clone | Vendor |
| --- | --- | --- |
| CCR7 PE | 150503 | R&D systems |
| CD3 CF-Blue | OC-515 | Immunostep |
| CD3 APC | OKT3 | Biolegend |
| CD3 FITC | OKT3 | Biolegend |
| CD4 APC | OKT4 | Biolegend |
| CD8 APC/Cy7 | HIT8a | Biolegend |
| CD8 PerCP | HIT8a | Biolegend |
| CD11c Pacific Blue | 3.9 | Biolegend |
| CD19 APC | HIB19 | Biolegend |
| CD40 FITC | 5C3 | Biolegend |
| CD45RO PE/Cy7 | UCHL1 | Biolegend |
| CD56 APC | 5.1H11 | Biolegend |
| CD107a APC | H4A3 | Biolegend |
| HIV-1 p24 PE | KC57-RD1 | Beckman coulter |
| HLA-DR PerCP | L243 | Biolegend |
| INFγ FITC | 4S.B3 | BD Pharmigen |
| PD1 PerCP | EH12.2H7 | Biolegend |
| TIGIT APC | A15153G | Biolegend |
| TIM3 FITC | F38-2E2 | Biolegend |
| Ghost Dye Red 780 |  | TONBO Bioscience |
| LIVE/DEA Fixable Yellow 405 |  | invitrogen |

### RNA extraction and qPCR mRNA quantification

Total RNA was isolated using the RNeasy Micro Kit (Qiagen) either from MDDCs from HIV-1 negative donors activated with individual or combined adjuvants, or from memory CD45RA-CD8+ T cells from PLWH and HIV-1 negative controls. cDNA was then synthesized using the reverse transcription kit from Promega. Transcriptional levels of PD-L1 (Forward: CACATCCATCATTCTCCCTTTTC, and Reverse: CATCCCAGAACTACCTCTGG), CD155 (Forward: TGGACGGCAAGAATGTGACC, and Reverse: ATCATAGCCAGAGATGGATACC) and Gal-9 (Forward: CACACATGCCTTTCCAGAAG, and Reverse: AAGAGGATCCCGTTCACCAT) on MDDC, and HIF1a (Forward: AGCCGAGGAAGAACTATGAACATAA, and Reverse: GTGGCCTGTGCAGTGCAA), GLUT1 (Forward: TCAACCGCAACGAGGAGAA, and Reverse: TCAACCGCAACGAGGAGAA), PDK1 (Forward: GTGGTTTATGTACCATCCCATCTCT, and Reverse: TCCATAGTGGCTCTCATTGCAT) and GAPDH (Forward: AGCCACATCGCTCAGACAC, and Reverse: GCCCAATACGACCAAATCC) on memory CD8 T cells were quantified by qRT-PCR using a final 0.5 μM concentration of each primer and SYBR Green GoTaq qPCR Master Mix (Promega). Amplification was performed in a StepOne Plus Real time PCR system (Applied Biosystems). Finally, relative mRNA expression was determined after normalization of each transcript to endogenous β-actin expression (Forward: CTGGAACGGTGAAGGTGACA, and reverse: CGGCCACATTGTGAACTT).

### Analysis of metabolic activity

Isolated CD8+ T cells from HIV negative donors or PLWH on ART at different times since treatment initiation were isolated as previously described and cultured for 2 h in RPMI 1640 media supplemented with 10 % Fetal Bovine Serum in the presence of IL-2 10 IU/ml. Then, 3×10^5^ cells from each donor were plated per replicate in DMED media adjusted to a pH of 7.4 in a p96 well plate (Seahorse XFe96 FluxPak Agilent Technologies 102416-100) in the presence of 0.125 mM Sodium pyruvate, 25 mM Glucose and 0.125 mM Glutamine. Cells were left alone or activated using Immunocult human CD3/CD28 T Cell Activator (Stem Cell Technologies) for 15 min; and 1 μM oligomycin, 1.5 μM CCCP, and 1 μM Rotenone plus 1 μM Antimycin A were sequentially injected following the Seahorse protocol specifications. Oxygen consumption rate (OCR) and extracellular acidification rate (ECAR) was measured using XF96 Extracellular Flux Analyzers (Seahorse); three measures (5 min) were obtained for each condition. Mitochondrial mass was evaluated as a control by flow cytometry using the MitoTracker Green staining probe (Invitrogen) at a concentration of 6.25 nM per million of cells. In some experiments, incorporation of glucose and fatty acid metabolism was determined by flow cytometry analysis of isolated CD8+ T cells from PLWH and control donors stained with 2-NBDG (Invitrogen) and 500/510 BODIPY (Molecular Probes) fluorescent probes, respectively following manufacture’s specifications.

### Statistics

Statistical significance of differences between the cells from different or within the same patient cohorts under different treatments were assessed using Mann Whitney U or Wilcoxon matched-pairs signed-rank tests. Multiple comparison correction using a Kruskal-Wallis test with post-hoc Dunn’s test correction method was applied when appropriate. Chi-square with Yate’s correction was used to compare differences in proportions of some parameters within different cell/patient populations. Non-parametric Spearman correlation was performed to test both individual correlations and to generate correlation networks. Statistical analyses were performed using GraphPad Prism 7.0 software. OCR and ECAR results were analyzed with Seahorse Wave software and then subjected to t-test multiple comparisons performed with GraphPad Prism 7.0 software and to a linear mixed model statistical analysis with R 4.1.1 version as previously described (36).

## RESULTS

### ART duration determines basal and DC-induced magnitude and polyfunctionality of HIV-specific CD8+ T cell in PLWH

For our study, we included a cohort of n=49 PLWH who had been under ART for at least 1 year and were characterized by undetectable HIV-1 plasma viremia in blood (Table 1). We first addressed intrinsic abilities of HIV-1-specific CD8+ T cells from these individuals to respond to Ag presentation by unstimulated and adjuvant-primed DC *in vitro*. To this end, we generated monocyte derived dendritic cells (MDDCs) from PBMCs from our cohort and cultured them in the presence of a specific pool of HIV-1 Gag peptides alone or in combination with the 2’3’-c’diAM(PS)2 STING agonist and the TLR3 ligand Poly I:C, as adjuvants. Higher level of maturation defined by up-regulation of CD40 was confirmed on adjuvant-treated MDDC from PLWH (**Supplementary figure 1 A, left panel**). We then evaluated the proportion of IFNγ+ CD8+ T cells from PLWH responding to HIV-1 Gag peptides presented by autologous MDDCs. Basal HIV-1 Gag specific-CD8+ T cell response was heterogeneous and two groups of PLWH were identified according to variations in the proportions IFNγ + cells above (Gag responders) or below (Gag non-responders) an optimal response cut-off of 2.5 fold-change from baseline (**Figure 1A)**. Interestingly, Gag responders were significantly enriched in individuals who had been under ART for more than a decade (from hereafter, LT-ARTp), while Gag non-responders were significantly enriched in PLWH on ART for less than 10 years (from hereafter, ST-ARTp) (**Figure 1B**). Significant differences in age were found in ST-ARTp and LT-ARTp groups, which were mainly composed by males (97% vs 67%, respectively). We did not find any other highly significant differences in other clinical parameters analyzed (Table 1). Next, we further assessed the impact of adjuvant-primed MDDCs loaded with HIV-1 Gag peptides on the magnitude and polyfunctionality of autologous HIV-1-specific CD8+ T cells responses in these two separate PLWH subgroups (**Supplementary figure 1B**). First, we confirmed that MDDCs from each patient subgroup were able to similarly and significantly increase CD40 expression after exposure to adjuvants (**Supplementary figure 1A, right**). In terms of magnitude, the induction of HIV-1 specific IFNγ+ CD8+ T cells from LT-ARTp compared to baseline did not further increase after co-culture with Gag adjuvant-primed MDDCs and responses were similar to those induced by Gag-MDDCs alone **(Figure 1C right panel)**. On the contrary, co-culture with Gag adjuvant-primed MDDCs significantly increased proportions of INFγ+ CD8+ T cells in ST-ARTp from baseline, in contrast to MDDC loaded with peptide only although levels did not reach those observed in LT-ARTp **(Figure 1C left panel)**. Importantly, Gag-loaded adjuvant-primed MDDCs significantly increased the frequencies of polyfunctional IFNγ+ CD107a+ CD8+ T cells in both ST-ARTp compared to baseline and in response to Gag only MDDCs **(Figure 1D, left)**. A similar significant increase on polyfunctionality of HIV-1 specific CD8+ T cells compared to baseline was observed in response to both Gag only and Gag-adjuvant primed MDDCs on LT-ARTp **(Figure 1D, right)**. Together, our results indicate that prolonged ART restores intrinsic abilities of HIV-specific CD8+ T cells to respond to DC treatment, while adjuvant stimulation enhances polyfunctionality of activated T cells.

**Figure 1.**
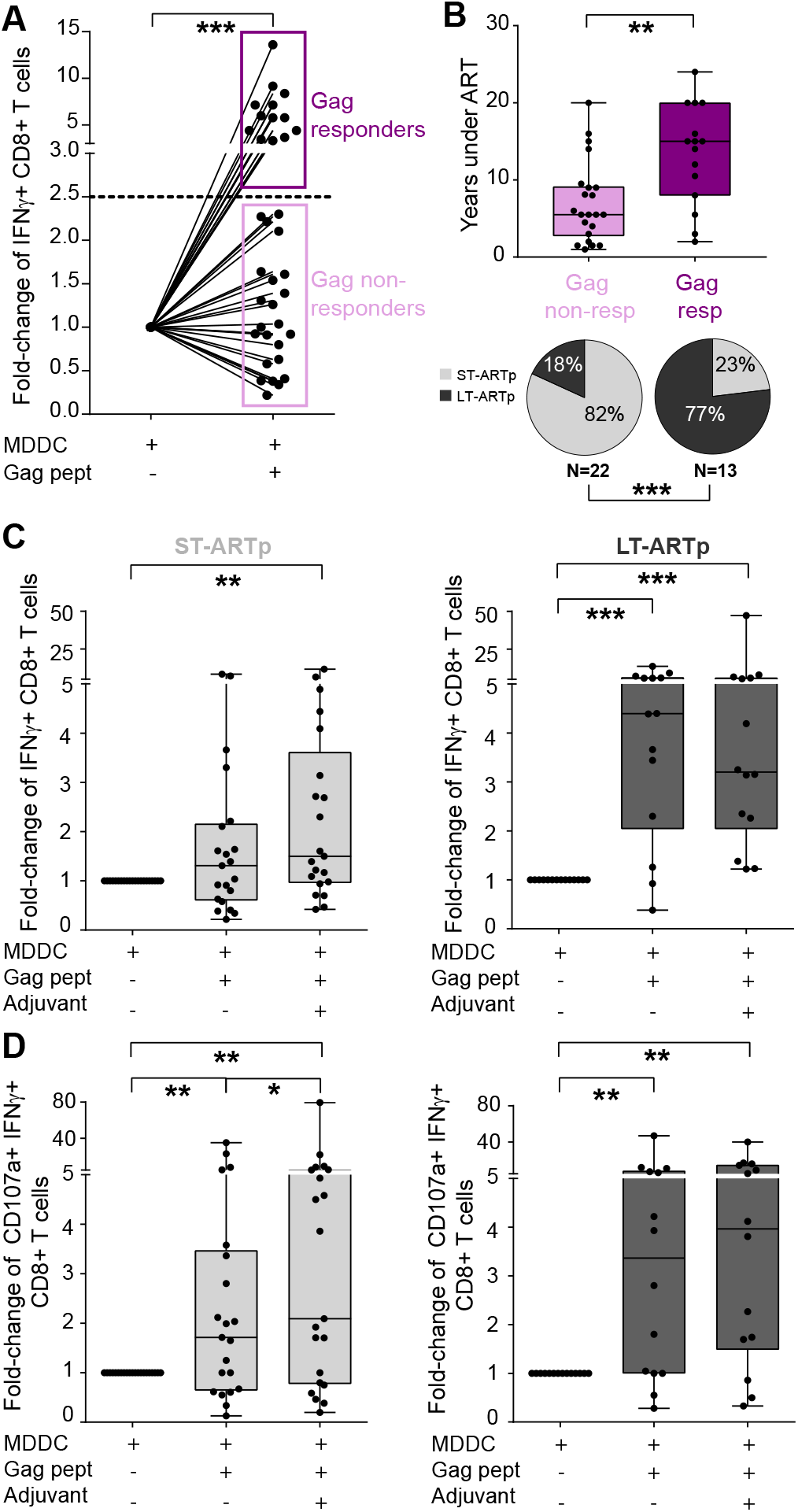
Magnitude and polyfunctionality of HIV-specific CD8+ T cell responses in PLWH on ART. (A): Fold-change in IFNγ expression from total live CD8+ T cells after Gag-peptide presentation by MDDCs, highlighting individuals who responded (Gag responders, in purple, fold-change > 2.5) and those who did not respond (Gag non-responders, in pink, fold-change <2.5) to basal HIV-1-Gag presentation by MDDC. (B): Upper panel showing whisker plots comparing years under ART from Gag responders (purple) and non-responders (pink); and lower panels showing pie-charts of the stratification between less than 10 years under ART (ST-ARTp, light grey), or equal or more than 10 years under ART (LT-ARTp, dark grey) included on each group of Gag responders or non-responders. Statistical significances were calculated using two-tailed Wilcoxon test (*p<0.05) and using a Chi-square test with Yates correction (***p<0.001), respectively. (C-D) Fold-change in IFNγ expression (C) and fold-change in polyfunctional responses (CD107a+ IFNγ+ cells) (D) from total live CD8+ T cells after stimulation of MDDCs in ST-ARTp (left plots, in light gray) and LT-ARTp (right plots, in dark gray). Statistical significance was calculated using two-tailed Wilcoxon test (*p<0.05; **p<0.01; ***p<0.001; ****p<0.0001).

### Differential restoration of cytotoxic function after DC treatment in CD8+ T cells from LT-ARTp and ST-ARTp

Next, we assessed whether changes on the magnitude and polyfunctionality of HIV-1 Gag-specific CD8+ T cell response after treatment with adjuvant-matured Gag-loaded MDDCs correlated with an increase in their cytotoxic capacity to eliminate HIV-1-infected CD4+ T cells. To address this, CD8+ T cells from ST-ARTp and LT-ARTp pre-stimulated with MDDCs primed in the absence or the presence of Gag peptides and adjuvants were subsequently co-cultured in bulk with autologous CD4+ T cells. These assays were performed in the presence of Romidepsin, a latency reversal agent, and the integrase inhibitor Raltegravir to prevent new rounds of infections in culture. After 24 h, frequencies of cells expressing intracellular HIV-1 p24 were evaluated by FACS (**Supplemental Figure 1C**). As shown in **Supplementary figure 2A**, Romidepsin treatment alone increased the frequencies of HIV-1 p24+ in CD4+ T cells from an important proportion of both ST-ARTp and LT-ARTp.

Notably, Gag-adjuvant-primed MDDCs enhanced cytotoxic capacities of CD8+ T cells to reduce frequencies of HIV-1 p24+ CD4+ T cells below baseline levels in 62% of the LT-ARTp individuals tested, while only 39% of the ST-ARTp exhibited these functional profiles (**Figure 2A-C**, in blue). On the other hand, CD8+ T cells that were unable to reduce proportions of HIV-1 p24+ CD4+ T cells after receiving MDDC stimulation were more represented among ST-ARTp (61%) compared to the LT-ARTp group (38%) (**Figure 2B-C**). Actually, in these functional assays from ST-ARTp we observed a significant increase in percentages of infected cells compared to baseline after exposure to Gag-adjuvant-primed MDDC, indicating the loss of antiviral properties in CD8+ T cells. In contrast, in the LT-ARTp displaying a dysfunctional phenotype after Gag-adjuvant-primed MDDC, the increase in the proportion of HIV p24+ CD4+ T cells detected compared to baseline was not even significant, suggesting that cells from these individuals retained some level of functionality (**Figure 2B-C**). Therefore, the improved phenotypic polyfunctionality of HIV-1-specific CD8+ T cells from PLWH induced in response to Gag-adjuvant-primed MDDCs is not always accompanied by increased cytotoxic properties of these cells, and years on ART seem to be a critical parameter that also determines recovery of effector function in CD8+ T cells.

**Figure 2.**
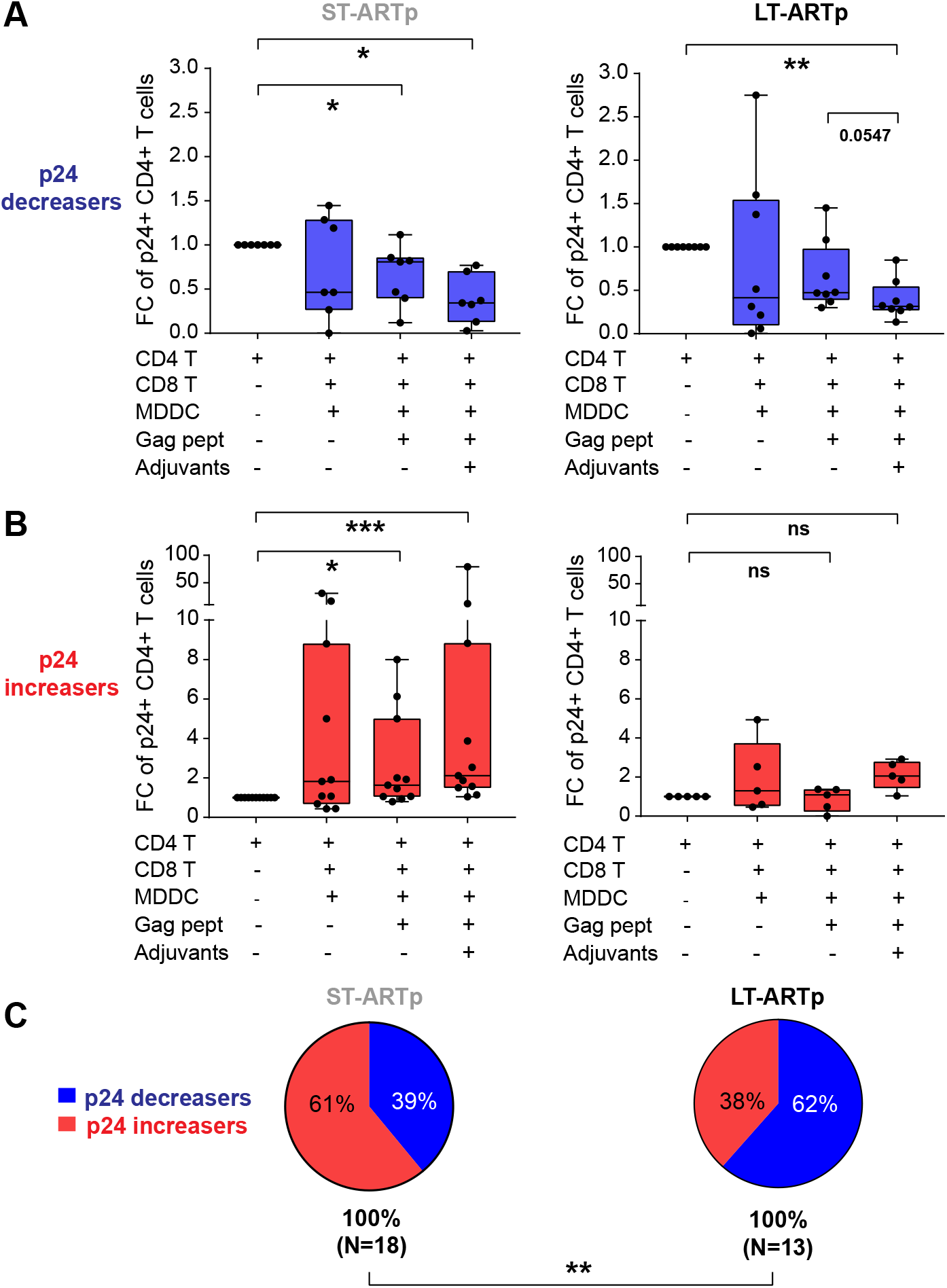
Cytotoxic function of CD8+ T cells from PLWH after DC-treatment. (A-B): Fold-change in intracellular HIV-1 p24+ cells from total live CD4+ T cells detected in the presence of autologous CD8+ T cells and primed with MDDC pre-cultured under the indicated conditions and relative to basal p24+ CD4+ T cell levels. Functional was stratified by reduction (A; p24 decreasers, in blue) or increase (B; p24 increasers, in red) of p24+ CD4+ T cell detection in ST-ARTp (left plots) and LT-ARTp (right plots) after MDDC treatment. Statistical significance was calculated using two-tailed Wilcoxon test (*p<0.05; **p<0.01). (C): Pie-charts showing percentage of increasers (red) and decreasers (blue) contained within the ST-ARTp subgroup (left) and the LT-ARTp subgroup (right). Pie-chart statistical significance of different was calculated using a Chi-square test with Yates correction.

### Differential memory subset distribution and distinct patterns of co-expression of checkpoint inhibitory receptors in CD8+ T cells from ST-ART and LT-ART groups

Differences between CD8+ T cells from ST-ARTp and LT-ARTp in their basal response to HIV-1 Gag-peptide stimulation and in the acquisition of cytotoxic function after treatment with Gag-adjuvant-primed MDDCs could be associated to either deficiency on DC activation, memory subset distribution or immune exhaustion of CD8+ T cells. We previously showed that adjuvant combination similarly increased the percentage of CD40+ MDDCs in both ST-ARTp and LT-ARTp, despite higher basal percentages of activation in ST-ARTp cells (**Supplemental Figure 1A**).

Since dysfunction of HIV-1-specific CD8+ T cell responses has been linked to reduced percentages of central memory (CM) cells (34), we next analyzed the proportions of different CD8+ T cell memory subsets in ST-ARTp and LT-ARTp. We observed that CD8+ T cells from LT-ARTp contained higher proportions of CCR7+ CD45RO+ CM cells, reaching similar percentages to HIV-1 negative controls (HC), and consistent lower frequencies of CCR7+ CD45RO-naïve cells (NA) (**Figure 3A**). On the contrary, CD8+ T cells from ST-ARTp were characterized by significantly lower frequencies of CM CD8+ T cells compared to LT-ARTp and tended to present higher proportions of CCR7-CD45RO+ effector memory (EM) and CCR7-CD45RO-terminally differentiated (TD) cells (**Figure 3A**). Preservation of CM cells was most evident in LT-ARTp that had received treatment shortly after diagnosis, but not in the case of ST-ARTp, highlighting the importance of both rapid ART initiation and adherence to preserve durable HIV-1-specific immunity (**Figure 3A, right plot**).

**Figure 3.**
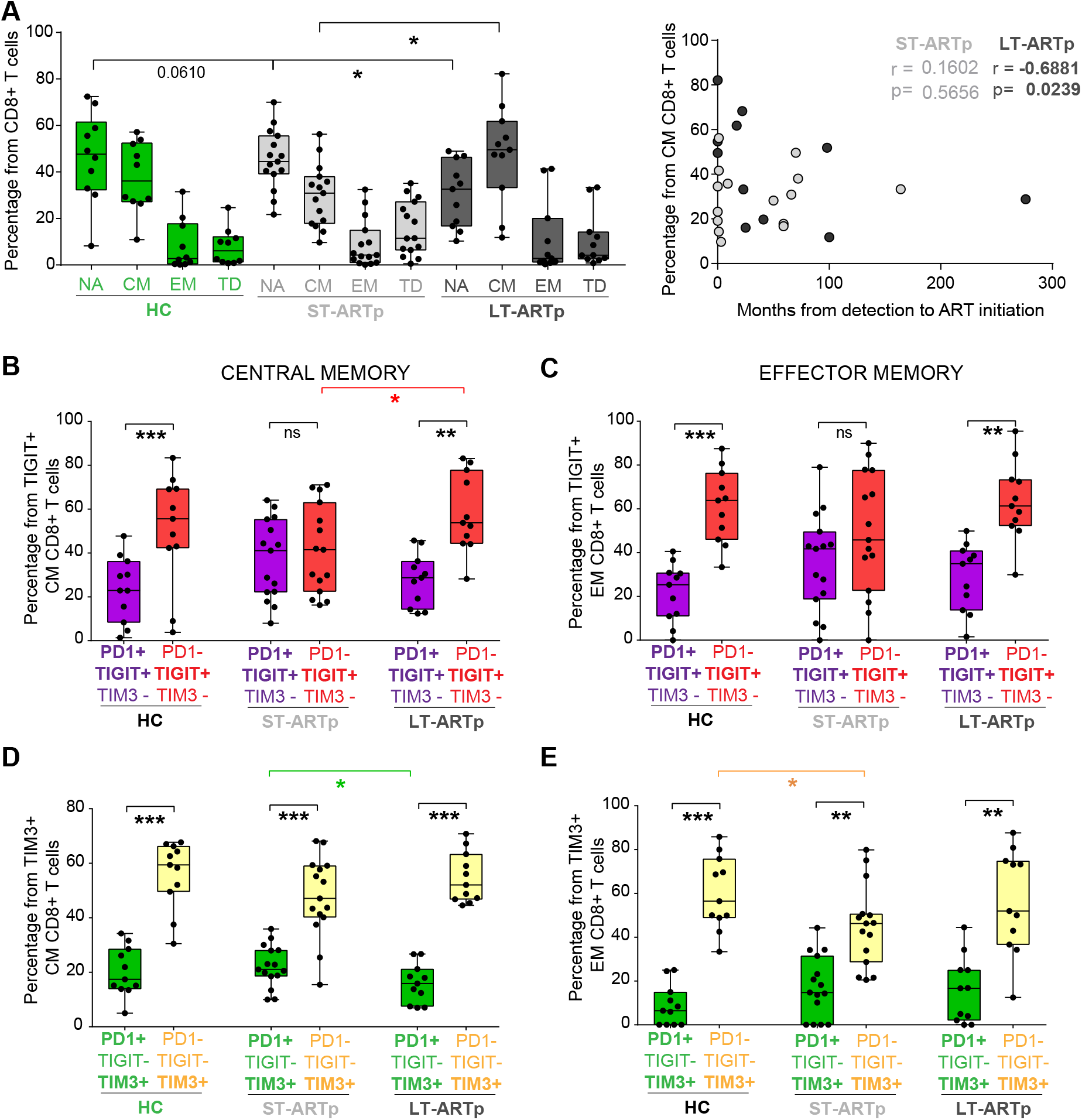
Memory subset distribution and co-expression of checkpoint inhibitor receptors in CD8+ T cells from different PLWH subgroups. (A): Percentage of naïve, NA (CCR7+ CD45Ro-); central memory, CM (CCR7+ CD45Ro+); effector memory, EM (CCR7-CD45Ro+); and terminally differentiated, TD (CCR7-CD45Ro-) from total live CD8+ T cells from HIV negative controls (HC; green bars), ST-ARTp (light gray bars) and LT-ARTp (dark gray bars). Statistical significance was calculated using two-tailed Mann-Whitney U test (*p<0.05). On the right, Spearman correlations between the percentage of CM CD8+ T cells and months since HIV-1 infection diagnosis to ART initiation for ST-ARTp (light gray) and LT-ARTp (dark gray dots). Spearman r and p values are shown on the left for each correlation. (B-E): Proportion of PD1+TIGIT+TIM3-(B-C; purple) or PD1+TIGIT-TIM3+ (D-E; green) populations from total TIGIT+ or TIM3+ cells, compared to PD1 - TIGIT+TIM3-(B-C; red) or PD1-TIGIT-TIM3+ (D-E; yellow) population from total TIGIT+ or TIM3+ populations. Statistical significance between the mentioned double and single positive populations within each participant group was calculated using two-tailed matched pairs Wilcoxon test (**p<0.01; ***p<0.001) and statistical significance between different individuals was calculated using two-tailed Mann-Whitney U test (*p<0.05).

Next, we asked whether differences of proportions in CD8+ T cell memory subsets from LT-ARTp and ST-ARTp could also be associated with the immune exhaustion state of these lymphocytes. In addition to inducing maturation, adjuvant treatment also increases expression of ligands for immunomodulatory checkpoint inhibitory receptors such as PD-L1, CD155 and Galectin-9 on MDDCs (**Supplementary figure 3A-C**), suggesting that bounding of these receptors might limit the ability of CD8+ T cells from PLWH to increase functional activities after MDDC stimulation. In addition, the co-expression of checkpoint inhibitory receptors has been associated to terminal exhaustion on T cells (22, 24, 37). Therefore, we determined the surface co-expression of PD1, TIGIT and TIM3 in CM and EM CD8+ T cells from ST-ARTp, LT-ARTp and HC by flow cytometry using boolean gating (**Supplementary figures 4A-B**). Notably, CD8+ T cells from ST-ARTp were significantly enriched on higher proportions of the PD1+ TIGIT+ TIM3-CM and EM cells in contrast to LT-ARTp, whose CD8+ T cells contained significantly higher proportions of the PD1-TIGIT+ TIM3-simple positive CM and EM cells, similarly to HC (**Figure 3 B-C, Supplementary figure 4A-B**). CD8+ T cells from ST-ARTp also showed higher co-expression of PD1 and TIM3 among the TIM3+ CM and EM population compared with the LT-ARTp (**Figure 3D-E, Supplementary figure 4A-B**). In addition, the proportions of PD1+ TIGIT+ TIM3+ cells were rare but significantly higher in EM CD8+ T cells from ST-ARTp, compared with LT-ARTp and HC (**Supplementary figure 4B**). Based on these results, we asked whether expression of these receptors is associated with memory T cell differentiation and/or proliferation. Interestingly, we observed an association between the differential dynamics of expression of PD1, TIGIT and TIM3, the proliferation and proportions of CM, and the recovery of CD8+ EM T cells in response to allogenic adjuvant-primed MDDC *in vitro* stimulation. Of note, proliferating CM and EM CD8+ T cells stimulated with MDDCs selectively induced TIM3 expression, whereas the induction of TIGIT and PD1 expression was weaker and not restricted to proliferating cells (**Supplementary figure 5A-C**). Therefore, our data indicate that non-overlapping co-expression profiles of different checkpoint inhibitory receptors induced after MDDC treatment are present on CD8+ T cells from PLWH at different times since ART duration.

### Differential induction of glycolytic metabolic activity in CD8+ T cells from ST-ART and LT-ART PLWH

Metabolic plasticity and glycolytic activity have been associated with exhaustion, effector function of CD8+ T cells and with their ability to mediate viral control (34, 38). Thus, we focused on studying the metabolic profiles of CD8+ T cells from LT-ARTp and ST-ARTp. To this end, we selected CD8+ T cells from LT-ARTp with high cytotoxic function and from ST-ARTp with dysfunctional characteristic and displaying opposite co-expression profiles of checkpoint receptors. These patients were also matched by age and sex with cells from HIV negative donors (**Tables 1 and 2**). Then, we studied mitochondrial respiration and the glycolytic rate by determining oxygen consumption (OCR) and extracellular acidification rate (ECAR) on CD8+ T cells from these PLWH groups and HC at baseline and after TCR/CD28 stimulation (**Figure 4A-B, Supplementary figure 6A-B**, also see Methods). While CD8+ T cells from HC induced significantly higher OCR after TCR stimulation, OCR values were significantly lower on stimulated CD8+ T cells from ST-ARTp, whereas cells from LT-ARTp mildly induced OCR upon TCR activation and differences were not significant compared to HC (**Figure 4A, Supplementary figure 7A, C**), suggesting potential defects on mitochondrial respiration on PLWH that were more marked in cells from ST-ARTp. In addition, we observed a significant increase of ECAR upon TCR activation in cells from all groups. Interestingly, CD8+ T lymphocytes from LT-ARTp showed increased ECAR induction, which was significantly higher than HC, suggesting higher ability to induce glycolysis (**Figure 4B, Supplementary figure 7B, D**). ECAR levels exhibited by CD8+ T cells from ST-ARTp after TCR stimulation were not significantly higher than those present in stimulated cells from HC (**Figure 4B, Supplementary figure 7B, D**). Importantly, we confirmed that the observed changes in the increase on ECAR in TCR-stimulated CD8+ T cells were not due to differences in glucose uptake or total mitochondrial mass between ST-ARTp, LT-ARTp and HC subgroups (**Supplementary figure 7E-F**). Alternatively, we observed that the basal transcriptional levels of the glucose metabolism regulator HIF1α was increased on memory CD8+ T cells from ST-ARTp compared to HC. In addition, mRNA levels of GLUT1 in memory CD8+ T cells from LT-ARTp were significantly lower compared to HC (**Figure 4 C**). The transcriptional levels of other molecules involved in the glycolytic metabolism such as PGK1 and GAPDH followed similar patterns as HIF1α on CD8+ T cells from ST-ARTp but were not significantly altered compared to HC (**Supplementary figure 8A**). Together, our results indicate that prolonged ART might preserve the ability of CD8+ T cells from PLWH to induce glycolysis and mitochondrial respiration upon TCR stimulation.

**Figure 4.**
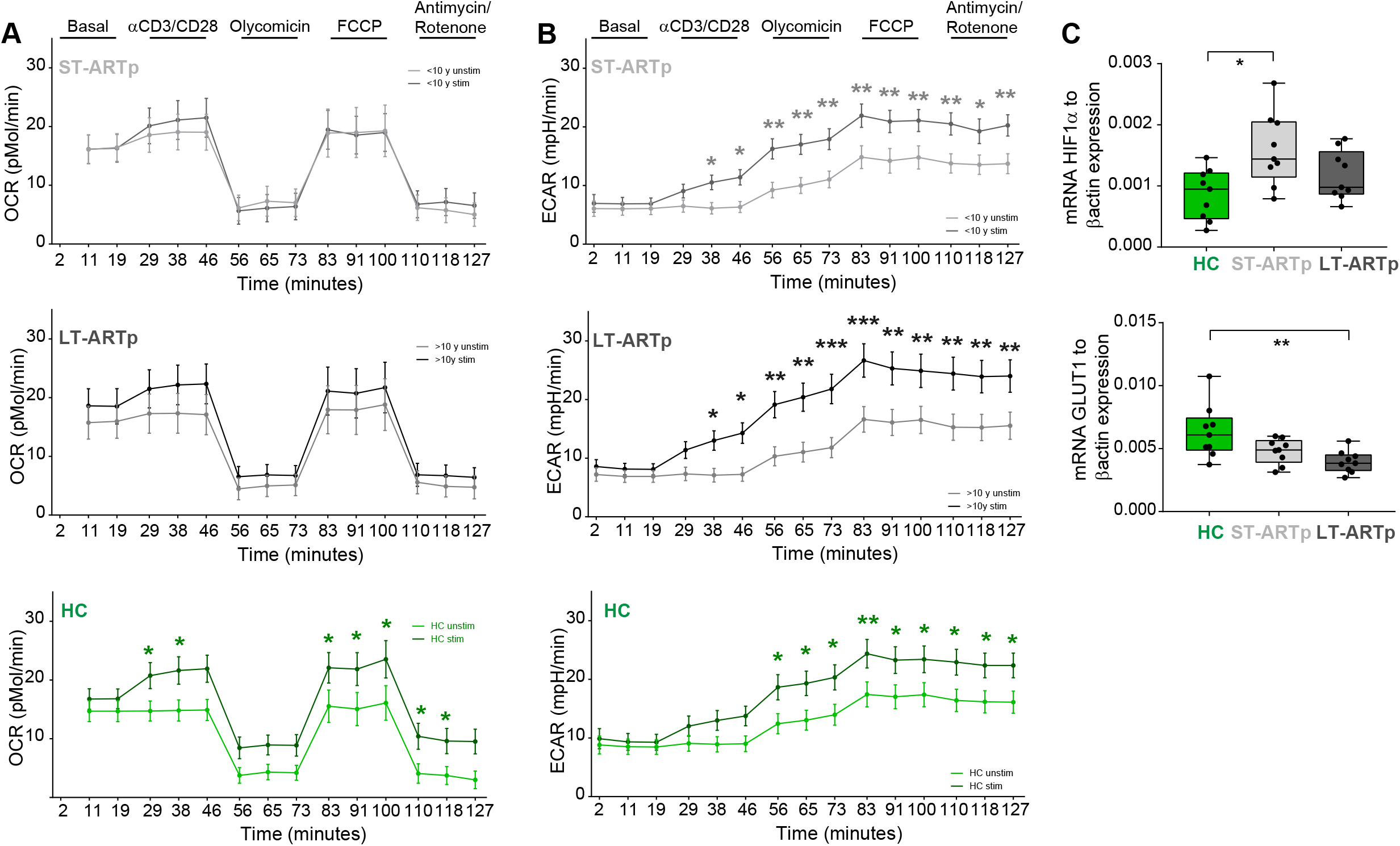
CD8+ T cell oxidative and glycolytic metabolism in PLWH. (A-B): Oxygen consumption rate (A; OCR) and extracellular acidification rate (B; ECAR) in CD8+ T cells from PLWH comparing basal (light line) and TCR activated (dark line) cells from ST-ARTp (upper plots; light gray), LT-ARTp (middle plots; dark gray), and HIV-1 negative controls (lower plots; green). Statistical significance per each timepoint was calculated with a multiple t test analysis. (C): RT-qPCR analysis of HIF1α (upper) and GLUT1 (lower) transcriptional expression normalized to β–actin mRNA levels in memory CD45RA-CD8+ T cells isolated from total PBMCs from HIV negative controls (HC; green) and PLWH (ST-ARTp; light gray and LT-ARTp; dark gray). Statistical significance between double and single positive populations was calculated using two-tailed Wilcoxon test (*p<0.05).

### Combined blockade of checkpoint inhibitory receptors and administration of glycolysis promoting drugs efficiently restores cytotoxic function of CD8+ T cell from ST-ARTp

PD1, TIGIT and TIM3 checkpoint inhibitory receptors can affect the glycolytic metabolism (28-30), but our results suggested that specific combinations of these receptors might differentially affect glycolysis induction and function of CD8+ T cells from ST-ARTp and LT-ARTp. Supporting this possibility, we observed significant positive correlations between the increase on glycolytic capacities, defined by delta ECAR from basal *versus* maximum values after TCR activation on CD8+ T cell, and the percentages of single positive TIGIT or TIM3 expression from the total CM or EM cells (**Figure 5A-B, upper panels; Supplementary figure 8B-C, upper panels**). In contrast, higher ratios of co-expression of PD1 and TIGIT or TIM3 compared with single expression of TIGIT or TIM3 in these memory CD8+ T cell subsets negatively correlated with the induction of glycolysis, respectively (**Figure 5 A-B, lower panels; Supplementary figure 8B-C, lower panels**). Therefore, our data demonstrate that specific patterns of TIGIT and PD1 co-expression in CM and EM CD8+ T cells correlate with impaired ability to induce glycolysis in total CD8+ T cells from PLWH.

**Figure 5.**
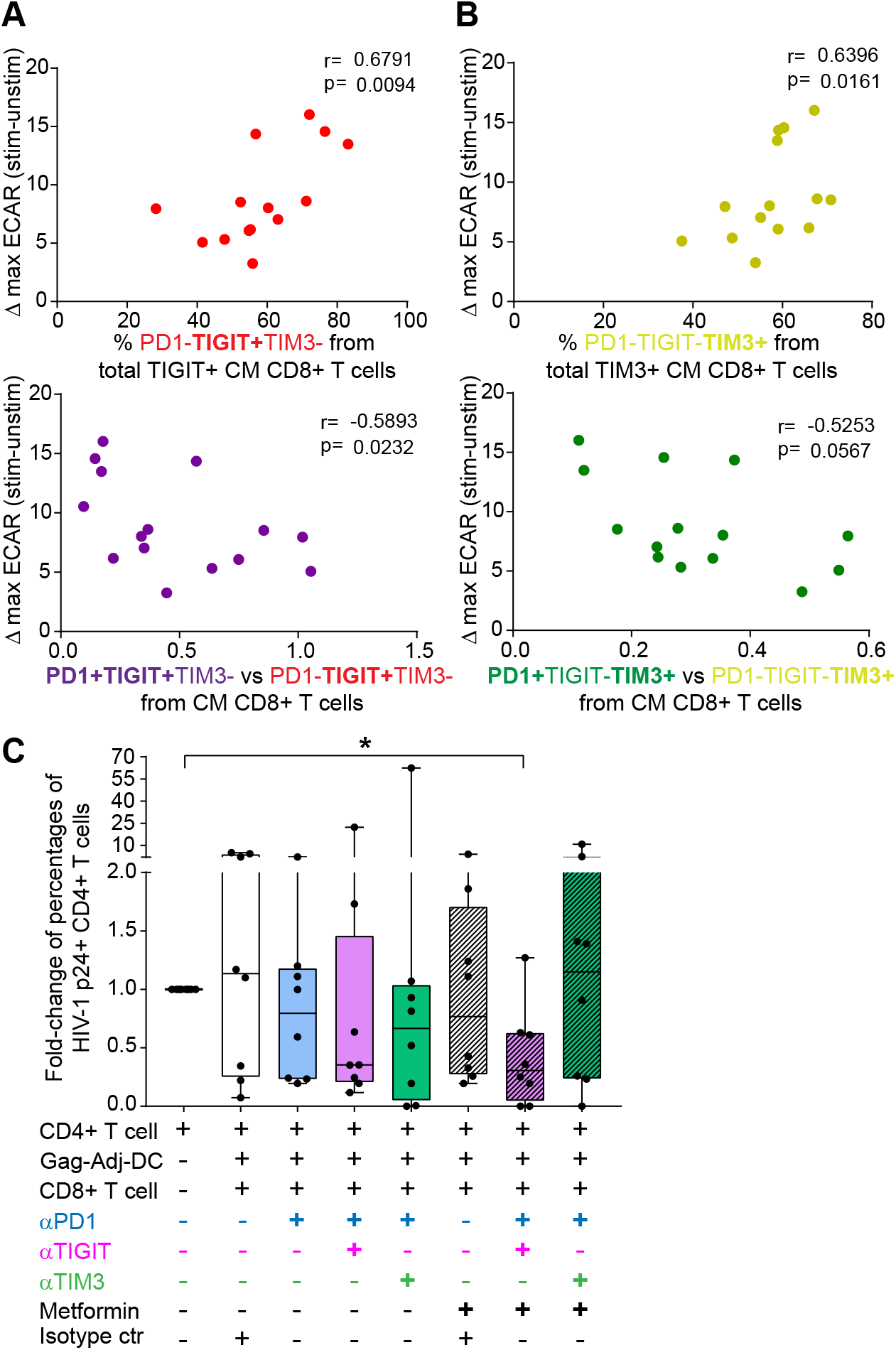
Correlation and fine-tuning of glycolytic metabolism and memory exhaustion in CD8+ T cell from chronic PLWH. (A-B): Spearman correlations between ΔECAR at maximal activation after TCR stimulation (73 minutes) and proportions of PD1-TIGIT+TIM3-(A, upper panel; red) or PD1-TIGIT-TIM3+ (B, upper panel; yellow) to total TIGIT+ or TIM3+ cells or ratios of the indicated populations (lower plots; PD1 *vs* TIGIT purple and red; PD1 *vs* TIM3 green and yellow) within central memory CD8+ T cells. Spearman r and p values are shown on the upper left area of each plot. (C): Fold-change in p24+ cells from total live CD4+ T cells from basal CD4+ T cells p24+ detection to the different MDDC-CD8+ T cell treated conditions. Light purple bars indicate the use of anti-PD1 and anti-TIGIT blocking antibodies in combination; light green bars indicate the use of anti-PD1 and anti-TIM3 blocking antibodies combination. Stripped bars indicate the use of 5 μM Metformin, either alone or in combination with the blocking antibodies. Statistical significance to the CD4+ T cell basal condition was calculated using two-tailed Wilcoxon test (*p<0.05).

Finally, we asked whether simultaneous blockade of multiple checkpoint inhibitors and/or treatment with pro-glycolytic drugs such as Metformin might represent potential modulatory treatments to more efficiently restore functional properties of CD8+ T cells from ST-ARTp after DC-treatment. To test these strategies, we stimulated dysfunctional CD8+ T cells from ST-ARTp with autologous adjuvant-primed and Gag-peptide loaded MDDCs in the presence of individual or a cocktail of antibodies specific for PD1, TIGIT and TIM3 alone or in combination with Metformin (see Methods). As shown in **Figure 5C**, combination of anti-TIGIT and anti-PD1 antibodies tended to increase the cytotoxic capacity of CD8+ T cells from ST-ARTp to eliminate p24+ CD4+ T cells in the presence of Romidepsin and Raltegavir, but it was not significantly different to CD4+ T cells cultured alone. Notably, the combined treatment of Metformin with anti-TIGIT and anti-PD1 mAbs was more efficient in restoring the cytotoxic capacities of CD8+ T cells from ST-ARTp and significantly reducing proportions of p24+ cells compared to CD4+ T cells alone (**Figure 5C**). Of note, corresponding isotype controls did not significantly alter proportions of HIV-1 p24+ cells detected. Dysfunction of CD8+ T cells was not corrected by individual blockade of PD1 or the combination of anti-PD1 and anti-TIM3 antibodies. Interestingly, the simultaneous blockade of TIM3 and PD1 in combination with Metformin seemed to even worsen their cytotoxic capacities, supporting a protective role of this receptor in line with our previous observations (**Figure 5C, Supplemental Figure 5**). Together, our data indicate that simultaneous blockade of TIGIT and PD1 combined with pro-glycolytic metabolism drugs can improve the functionality of CD8+ T cells from ST-ARTp after MDDC stimulation.

## DISCUSSION

The present study evaluated the efficacy of adjuvant-primed MDDC restoring cytotoxic capacities of CD8+ T cell from different groups of PLWH, who are under ART and characterized by undetectable plasma HIV-1 viral load, as a potential therapeutic candidate to treat HIV infected individuals. Previous studies using adjuvants have suggested that targeting innate immunity could be useful to potentiate HIV-1-specific CD8+ T cells. In this sense, innate-specific adjuvant-mediated activation displayed promising results when CD40+ cells were activated through TLR3 (39). However, these approaches failed in preventing viral rebound after stopping ART, probably because adjuvants were administered systemically and could have induced a general hyper-inflammatory response favoring HIV-1 replication (40-45). Our therapeutic approach is based on the activation of DCs with the combined use of TLR3 and STING agonists in the presence of a pool of HIV-1-Gag peptides, which has been previously associated with increased polyfunctional HIV-1 specific CD8+ T cell responses in lymphoid tissues *in vivo* (19). Compared with previously published approaches, our strategy allows avoiding bystander activation of other cells than DCs in response to adjuvants. In this study, we have shown that this method can also be useful to induce polyfunctional HIV-1-specific CD8+ T cell from PLWH responses resulting in reduction of HIV-infected CD4+ T cells *in vitro*.

We have also demonstrated that the years under antiretroviral treatment determine the basal capacity of HIV-1 specific CD8+ T cells to respond to MDDC stimulation and antigen presentation. We have observed differences on checkpoint inhibitory receptor phenotypes and on frequencies of central and effector memory CD8+ T cells that are associated with the time since ART initiation. These results are in line with previous studies reporting the recovery from immune exhaustion upon HIV-1 pharmacological suppression, as well as differences in metabolism in these treated individuals (34, 46-48). Our study identifies for the first time immunological and metabolic patterns that are specifically associated with effective responses of CD8+T cells to DC-based treatment in different subgroups of PLWH.

We have defined that prolonged treatment duration is required to restore phenotypical and effector capacities of CD8+ T cells and to respond to MDDC-based HIV-1 vaccines. Those results are in line with previous studies by Perdomo-Celis *et al* (49), demonstrating that CD8+ T cell cytotoxic and polyfunctional capacities are reduced in PLWH compared to HIV negative individuals, and that ART only slightly reinvigorates the cytolytic capacities after two years of treatment. We have found a negative correlation between the time from HIV diagnosis to ART initiation and preserved proportions of CM CD8+ T cell subpopulation in LT-ARTp. These findings are in agreement with previous studies suggesting that early ART initiation preserves immune function of memory T cells (50). One of the limitations of the present study is that we considered IFNγ and CD107a co-expression as a readout of polyfunctional HIV-1-specific T cells without taking into account their co-expression with TNFα, which also has been associated with durable and efficient CD8+ T cell responses in HIV-1 EC and in humanized mice (19, 51). We neither assessed whether detected polyfunctional CD8+ T cells completely corresponded the antigen-specific cells induced by our adjuvant primed MDDC-based therapy by analyzing TCR specificity by tetramer staining. Thus, further studies evaluating the HIV-1-specific TCR CD8+ T cell responses will be required to further confirm our results.

On the other hand, our study provides novel and relevant information about the parameters determining the exhaustion and metabolic state of CD8+ T cells and their connection with the response to DC therapy. We have described that ST-ARTp contain lower proportions of CM CD8+ T cells and that those cells are characterized by higher co-expression of TIGIT and PD1. Interestingly, LT-ARTp displayed higher percentages of CM CD8+ T cells, which were more enriched by single expression of TIGIT or TIM3. While previous studies had described that co-expression of checkpoint inhibitory receptors might be associated with reduced cytotoxic activity of HIV-1 specific CD8+ T cells in PLWH (37), our study provides new information suggesting that different combinations of specific checkpoint inhibitory receptors might differentially affect functional exhaustion of T cells. The role of PD1 limiting the development and effector function of memory CD8+ T cells, and the beneficial effect of PD1 blockade have already been described in cancer and also in other viral infections. Anti-PD1 mAb based therapies have yielded promising results in animal models of SIV-infection in macaques, enhancing specific CD8+ T cell responses and reducing the plasma viral load, resulting in enhanced survival of the animals. However, they were not sufficient to prevent viral rebound after treatment interruption (15, 52, 53). Thus, modulation of additional checkpoint inhibitory receptors and metabolic pathways might be required to more efficiently reinvigorate functional HIV-specific CD8+ T cells in PLWH. In this regard, we have demonstrated that the combined blockade of anti-PD1 and anti-TIGIT improves the cytotoxic capacities of CD8+ T cells from ST-ARTp against HIV-infected CD4+ T cells more effectively than anti-PD1 blockade alone. These data are supported by similar results in a gastric cancer animal model (29). Our results also indicate that combined TIM3 and PD-1 blockade seemed to disrupt cytotoxic function of CD8+ T cell from ST-ARTp individuals. Although TIM3 blockade inhibited T regulatory function and has yielded promising results on different cancer model, TIM3 has also been reported as a marker of proliferating IFNγ+ Th1 cells, key in the antiviral response. The effector function of these cells could be dependent on TIM3 expression (54, 55). In this regard, CD8+ T cells co-expressing PD1 and TIM3 are more prone to respond to PD1 blockade in cancer therapies (33). Therefore, it is unclear whether TIM3 expression might be beneficial or detrimental for CD8+ T cell response generation in PLWH and further studies are required to address this question.

Another key aspect of our study is the fact that we have analyzed the association between phenotypical and functional differences on CD8+ T cells from HIV-1 chronic patients exposed to DC with checkpoint inhibitory markers expression and other processes associated to exhaustion, such as mitochondrial respiration and the glucose metabolism. Previous studies have reported a correlation between glycolysis increase and effective cytotoxic functions of CD8+ T cells against HIV-infected cells in treated PLWH (56). Reduced glycolysis has been associated to HIV-1 latency and oxidative stress in infected individuals (57). We reported higher induction of ECAR after TCR stimulation in CD8+ T cells from LT-ARTp displaying effective cytotoxic function against HIV-1-infected cells, whilst enrichment of TIGIT+ PD1+ in CD8+ T cells from ST-ARTp associated to reduced ability to increase ECAR upon TCR activation. We were able to describe a correlation between CD8+ T cell effector and cytotoxic functions upon MDDC activation, basal CD8+ T cell memory exhaustion phenotypes, metabolic dysfunctional state and years under ART treatment in a cohort of PLWH on ART. However, an important limitation from our study is that we did not directly address the metabolic and functional properties of CD8+ T cells from ST-ARTp and LT-ARTp isolated based on the differential expression of checkpoint receptors. In addition, immunosenescence has been described as a process affecting metabolism thorough adult life to elderly, but we have not extensively analyzed the impact of differences in age on our study cohorts by excluding the youngest and oldest individuals. Oxidative metabolism seems to be largely altered in PLWH, however, the mechanisms leading to these observations have not been assessed. The role of T cell factor-1, which is expressed by effective long-lived PD1 low memory CD8+ T cells during chronic infections (33, 58), and mitochondrial fission and fusion, which also play a critical role on effector T cell function and memory maintenance (59, 60) may be also playing a role in these processes, and will be of interest to fully understand the mechanisms driving the differences on the metabolic dysfunction described in this study.

Our PLWH cohort was defined by individuals who were under ART, with undetectable plasma HIV-1 viral load (<20 mRNA copies/ml), with no co-infection with HCV, and CD4+ T cell counts higher than 400 cells/ml. In this regard, we focused on PLWH who more likely contain preserved memory CD8+ T cells able to mediate competent immune responses, as reduction of antigen exposure on treated PLWH has been described to partially restore CD8+ T cell function and reduce exhaustion, compared to non-treated PLWH (61-63). This allowed us to evaluate the capacity of our adjuvant-DC strategy to induce CD8+ T cell responses in PLWH at different times since treatment initiation. However, some of the observed responses and differences might not be present in viremic PLWH with low CD4+ and CD8+ T cell counts. In these patients, memory CD8+ T cell populations could be diminished and balanced to a more terminally and exhausted memory phenotype, and therefore less capable of mediating competent cytotoxic effector responses even after DC stimulation. Therefore, further analysis should address the effectiveness of DC therapy and the proposed combined treatment with blocking antibodies and glycolysis inductors for this particular populations of PLWH reinvigorating their memory T cells. In conclusion, our study identifies specific immunometabolic parameters for different groups of PLWH defined by anti-retroviral treatment duration, non-overlapping expression of checkpoint inhibitory receptors and metabolic state of CD8+ T cells that can be modulated through personalized therapies to fine-tune functional HIV-1 CD8+ T cell responses, providing new tools to advance and improve DC-based HIV-1 vaccines.

## Acknowledgements

We would like to thank the NIH AIDS Reagent Program, Division of AIDS, NIAID, NIH for providing HIV-1 PTE Gag Peptide Pool from NIAID, DAIDS (cat #11057) for the study. We would also like to thank Álvaro Serrano Navarro for his help on adapting the linear mixed model previously described by Martin-Cofreces N. *et al* (35) to our data.

## Author contributions

EMG developed the research idea and study concept, designed and supervised the study.

MCM designed and conducted most experiments of the study.

ISC contributed to functional assays.

IT, MCM, CDA performed analysis of checkpoint receptors and proliferation in MLR experiments from the study.

NMC, MCM and EMG designed and performed Seahorse experiments.

MCM, ISC, CDA processed peripheral blood samples from PLWH and HIV negative donors.

MJB, HDF, NMC provided critical feedback during experimental design and execution phases of the studies and were directly involved in the experiments.

MJC provided reagents for transcriptional analysis of metabolic regulators and provided critical feedback.

FSM, AA, MAMF, IDS, LGF, and JS provided peripheral blood from PLWH and HIV negative controls, reagents, clinical expertise and participated on the analysis of the data.

## Data Availability

The data that support the findings of this study will be available upon reasonable request to the corresponding author of the study.

## SUPPLEMENTARY FIGURE LEGENDS

**Supplementary figure 1.**
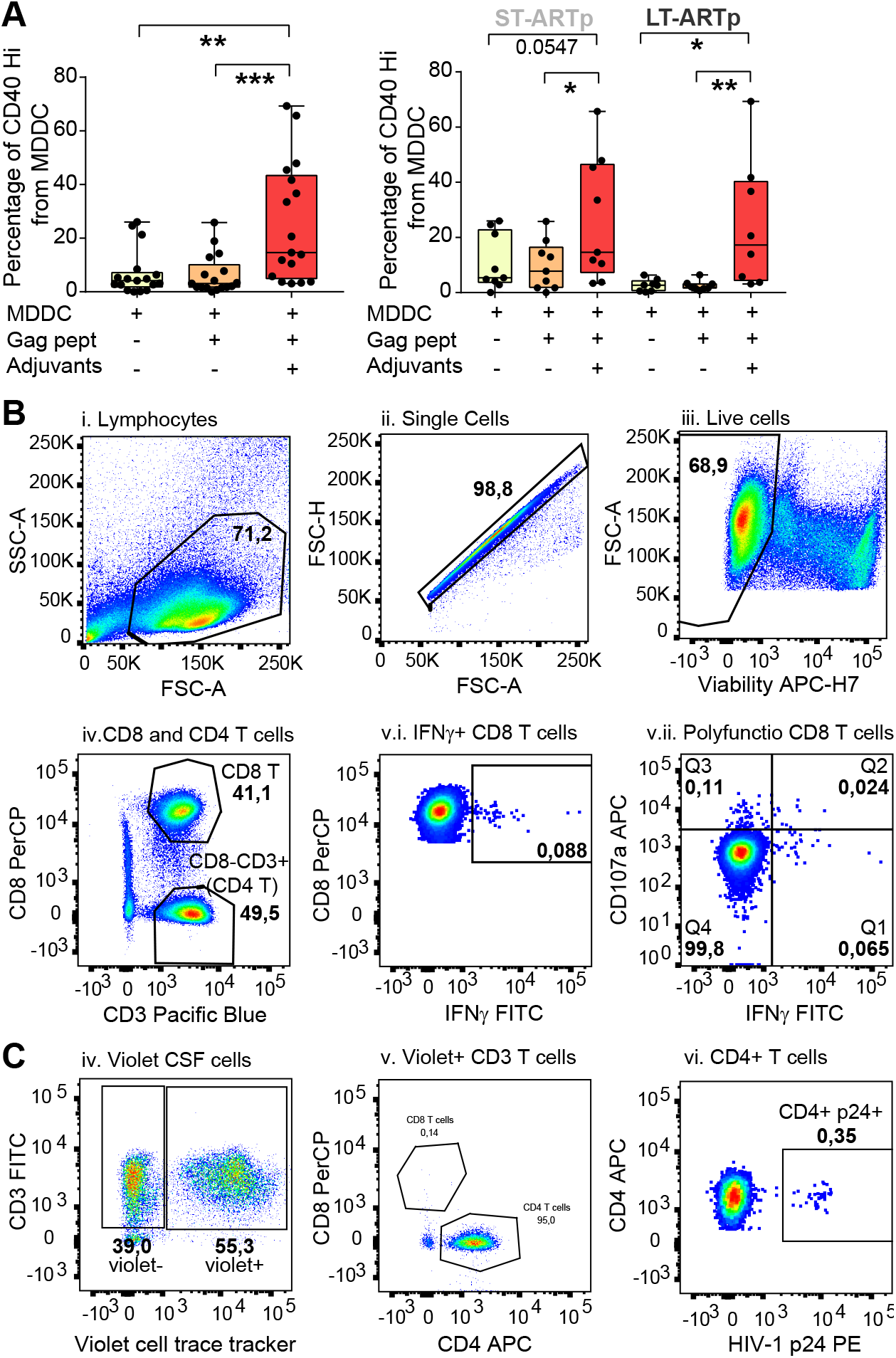
Analysis of MDDC activation and flow-cytometry gating strategies identifying HIV-specific CD8+ T cell responses and p24+ CD4+ T cell in PLWH. (A): Quantification of proportions of CD40Hi MDDCs generated from total PLWH (left) or individual ST-ARTp (left) and LT-ARTp (right) groups cultured for 16 h in media alone or in the presence of a pool of HIV-1 Gag peptides in the absence or the presence of Poly I:C and 2’3’-c’diAM(PS)2 adjuvant combination. Significant statistical differences after treatment were calculated using two-tailed matched pairs Wilcoxon test (*p<0.05; **p<0.01). (B,C): Representative flow cytometry dot plots defining gating strategy for (B) IFNγ+ CD8+ T cells (v.i.) and polyfunctional (IFNγ+ CD107a+) (v.ii.) CD8+ T cells identified from total single viable CD3+CD8+ T lymphocytes and for (C): HIV-1 p24+ CD4+ T cell identified from total violet CSF stained CD3+CD4+T cell.

**Supplementary figure 2.**
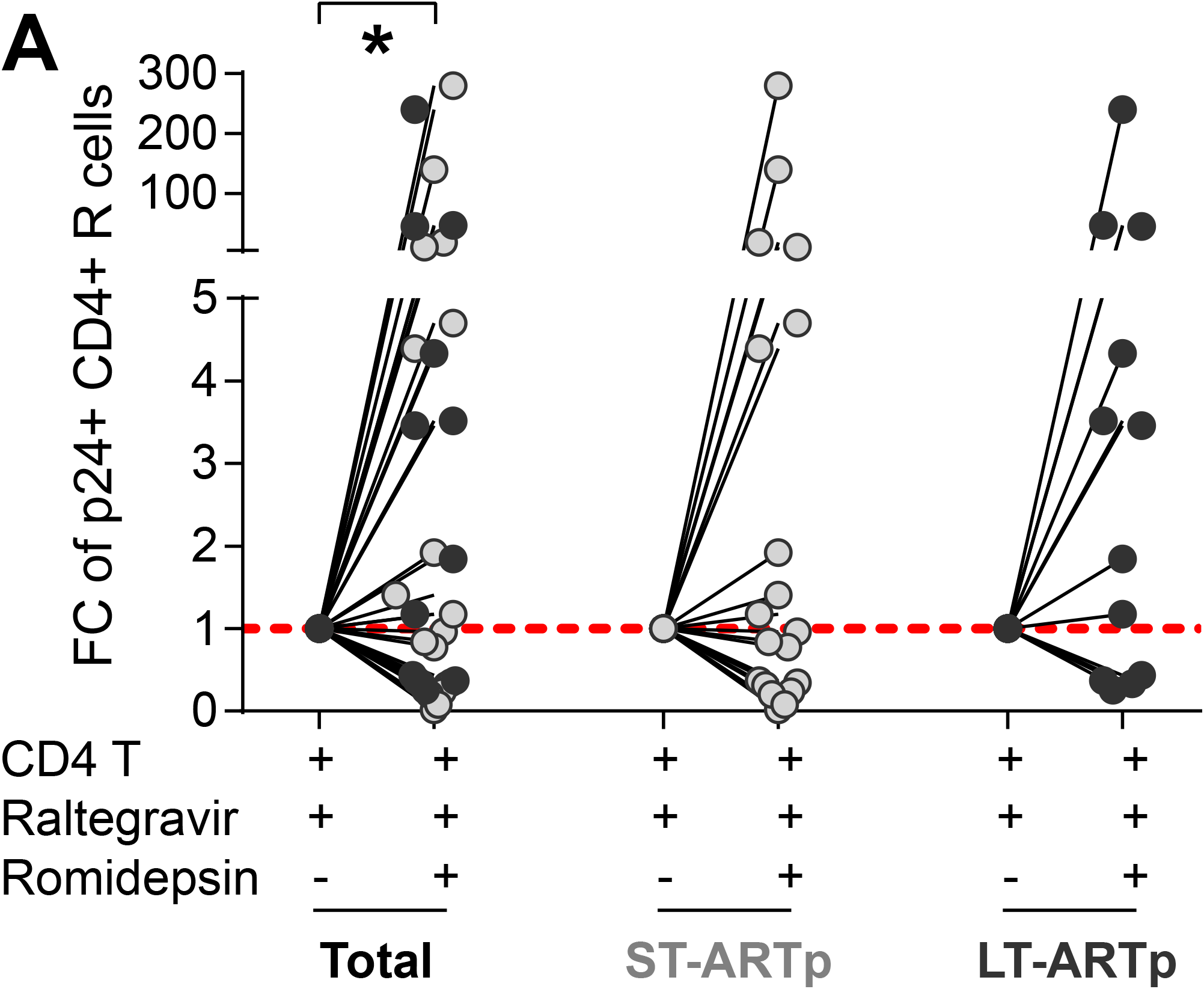
HIV-1 p24+ CD4+ T cells from PWLH induced after Romidepsin treatment. (A): Fold-change in intracellular HIV-1 p24+ cells from total live CD4+ T cells treated with Romidepsin and Raltegavir relative to basal detection of p24 on CD4+ T cells treated with Raltegavir alone. Left plot shows all PWLH combined, middle plot shows ST-ARTp and right plot shows LT-ARTp. Significant statistical differences after treatment were calculated using two-tailed matched pairs Wilcoxon test (*p<0.05).

**Supplementary figure 3.**
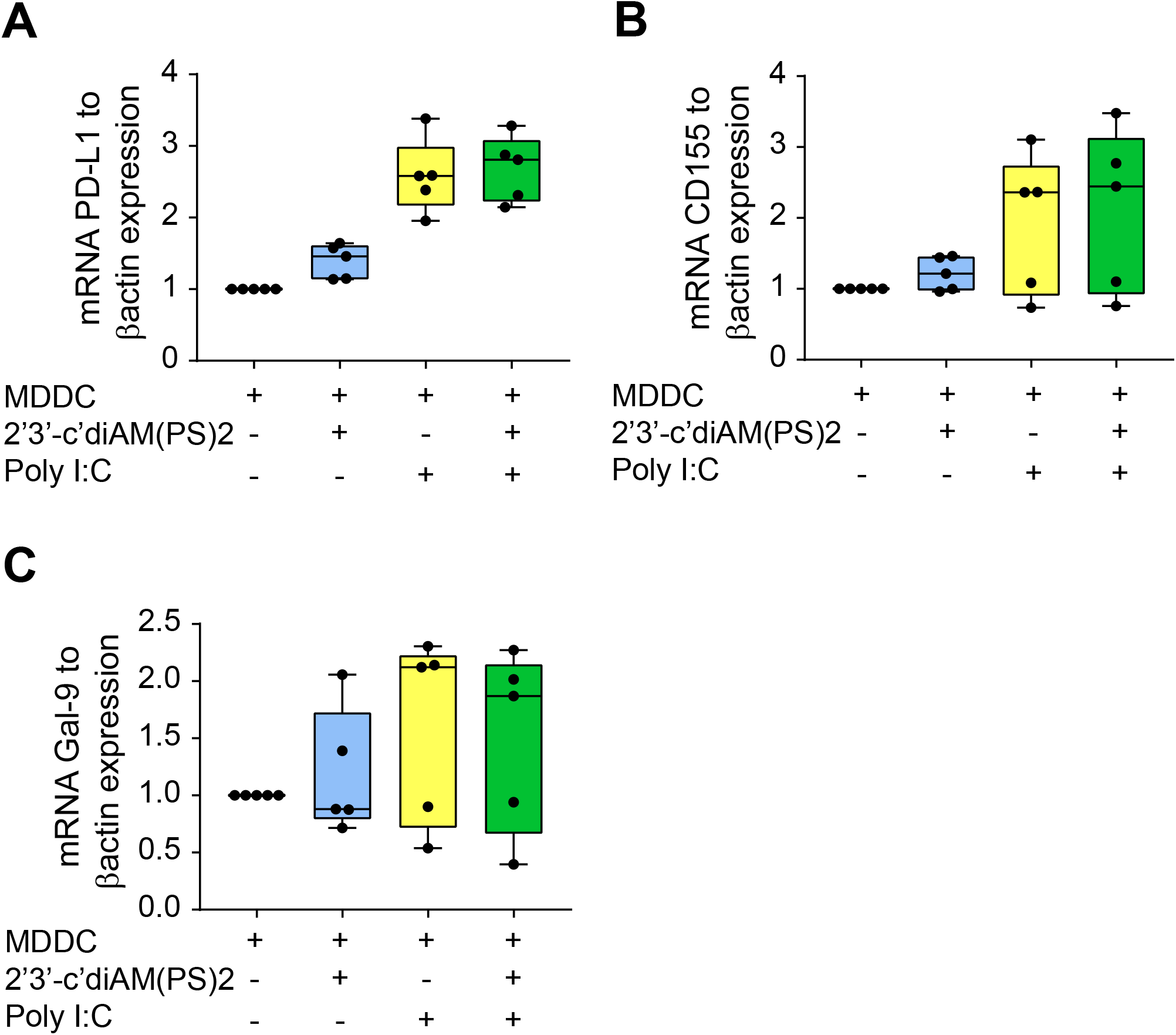
Impact of adjuvant-maturation on expression of checkpoint inhibitory ligands in MDDC. (A,B,C): RT-qPCR analysis of PD-L1 (A), CD155 (B) and Galectin 9 (Gal-9,C) transcriptional expression normalized to β–actin mRNA levels in MDDCs generated from HIV-1 negative controls and cultured for 16 hours with media alone or in the presence of either individual or combined STING and TLR3 agonists (n=5 independent experiments). Statistical significance between double and single positive populations was calculated using two-tailed Wilcoxon test (*p<0.05).

**Supplementary figure 4.**
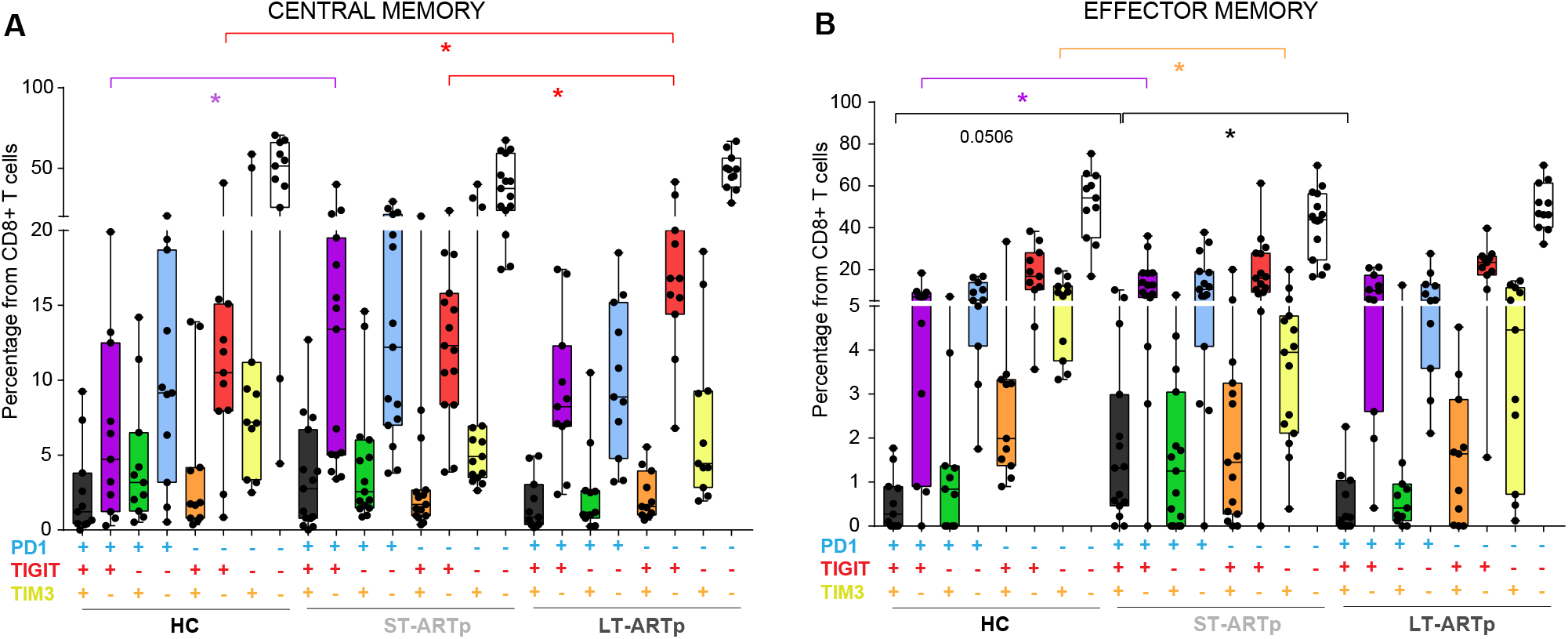
Expression of checkpoint inhibitory receptors in different CD8+ T cell memory subsets. (A-B): Boolean gating of PD1, TIGIT and TIM3 co-expression combinations on either central memory (A) or effector memory (B) CD8+ T cells. Statistical significance between groups was calculated using two-tailed Mann-Whitney U test (*p<0.05). For clarity purposes, significances are highlighted on the color of each corresponding population.

**Supplementary figure 5.**
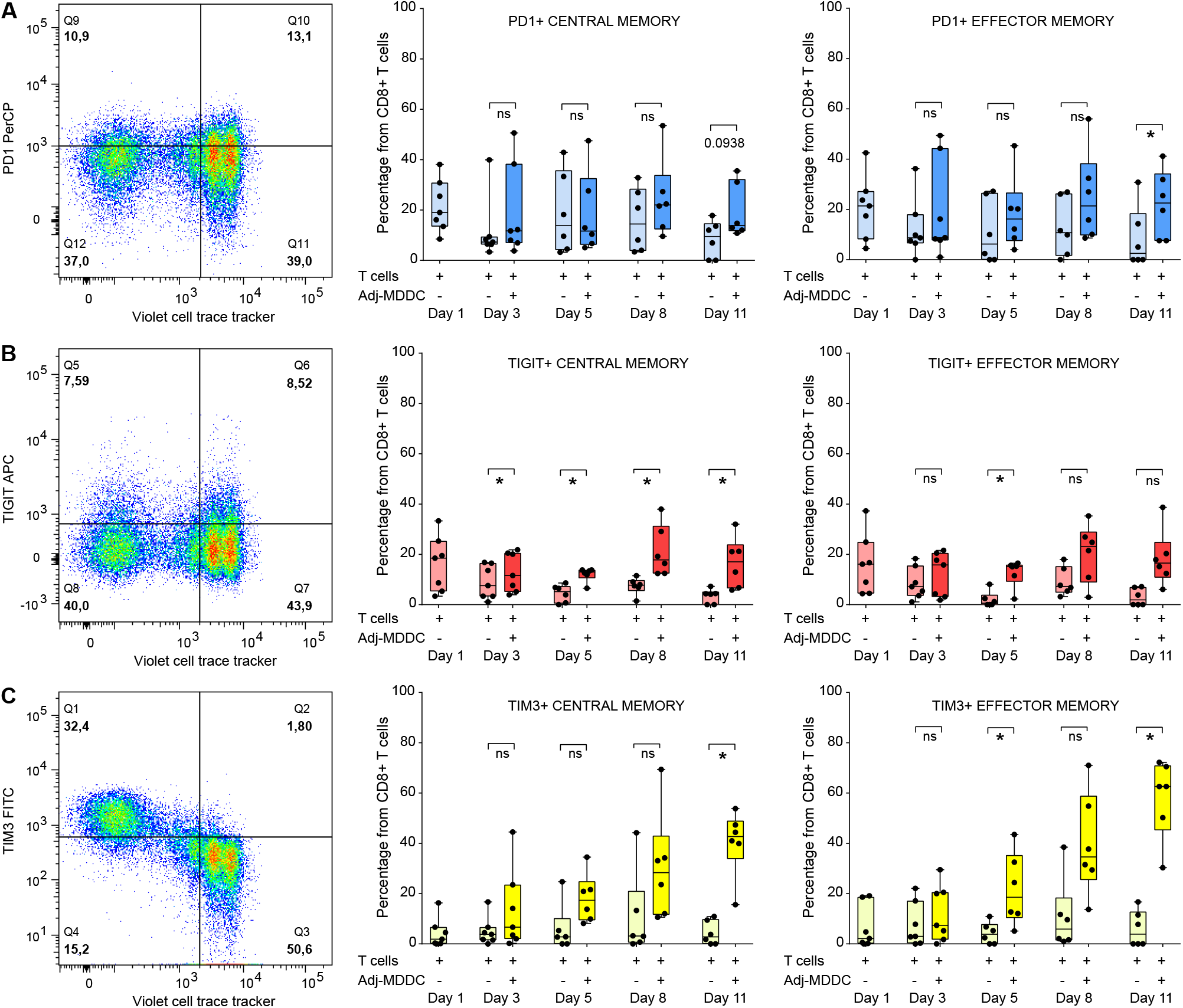
Dynamics of checkpoint inhibitory receptors expression on memory CD8+ T cell subsets cultured with MDDC. (A-C): Time-course of proportions of PD1+ (A), TIGIT+ (B) and TIM3+ (C) cells on central (middle plots) and effector (right plots) memory CD8+ T cells from total T cells isolated from HIV negative donors cultured for 11 days in the presence or absence of adjuvant-activated allogenic MDDCs. Representative flow cytometry dot plots showing proliferation (violet CSF+) *vs* expression of each of the checkpoint inhibitory receptors analyzed on central memory CD8+ T cells after 8 days of culture are shown on the left show.

**Supplementary figure 6.**
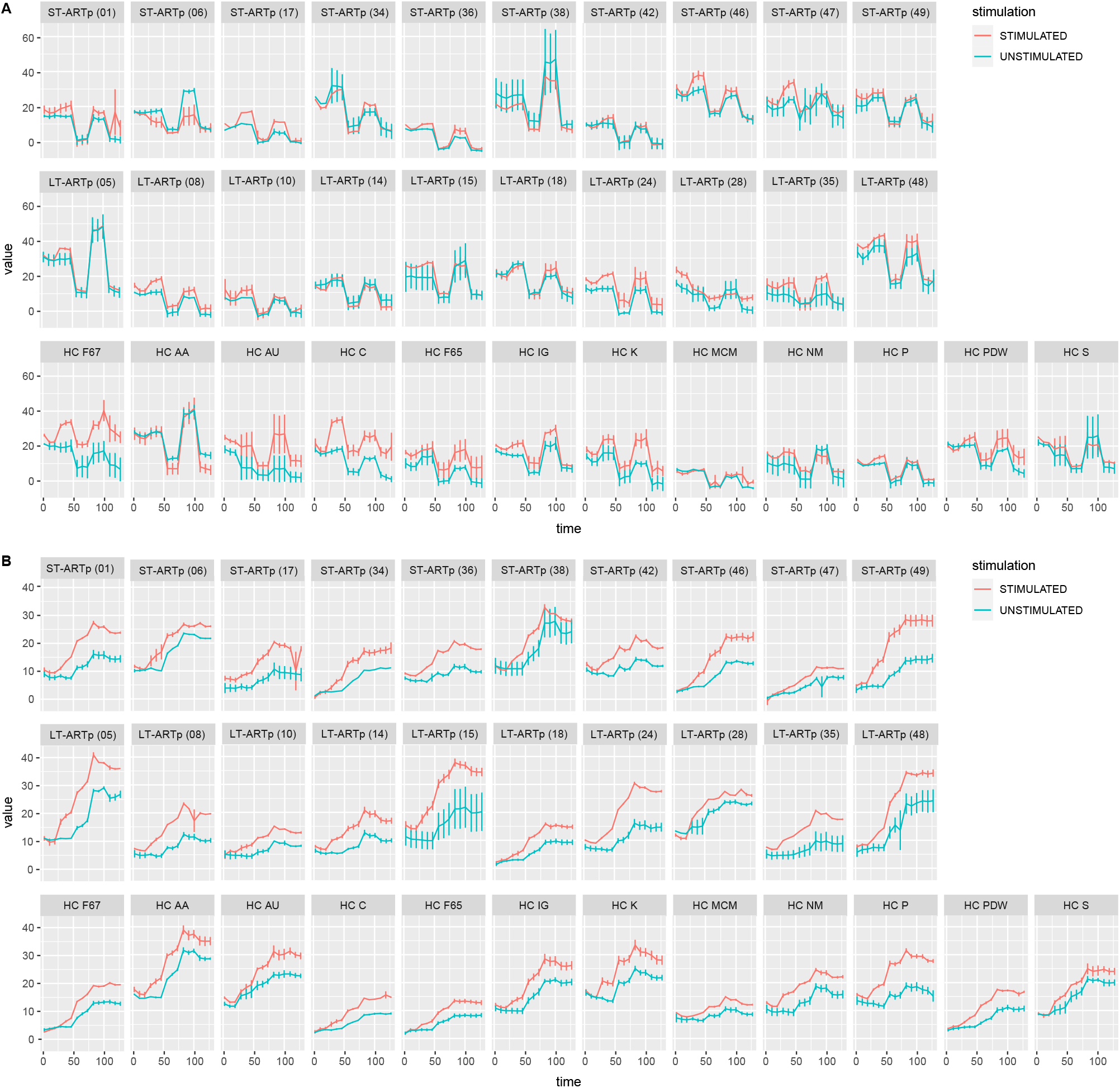
OCR and ECAR values for each individual patient analyzed. (A-B): Individual plots showing OCR (A) and ECAR (B) mean and standard error for basal and activated CD8+ T cells per timepoint from each study participant (ST-ARTp upper rows; LT-ARTp middle rows; HIV negative bottom rows) used for the pooled analysis. Means and standard errors were obtained with R 4.1.1 version dplyr and ggplot packages.

**Supplementary figure 7.**
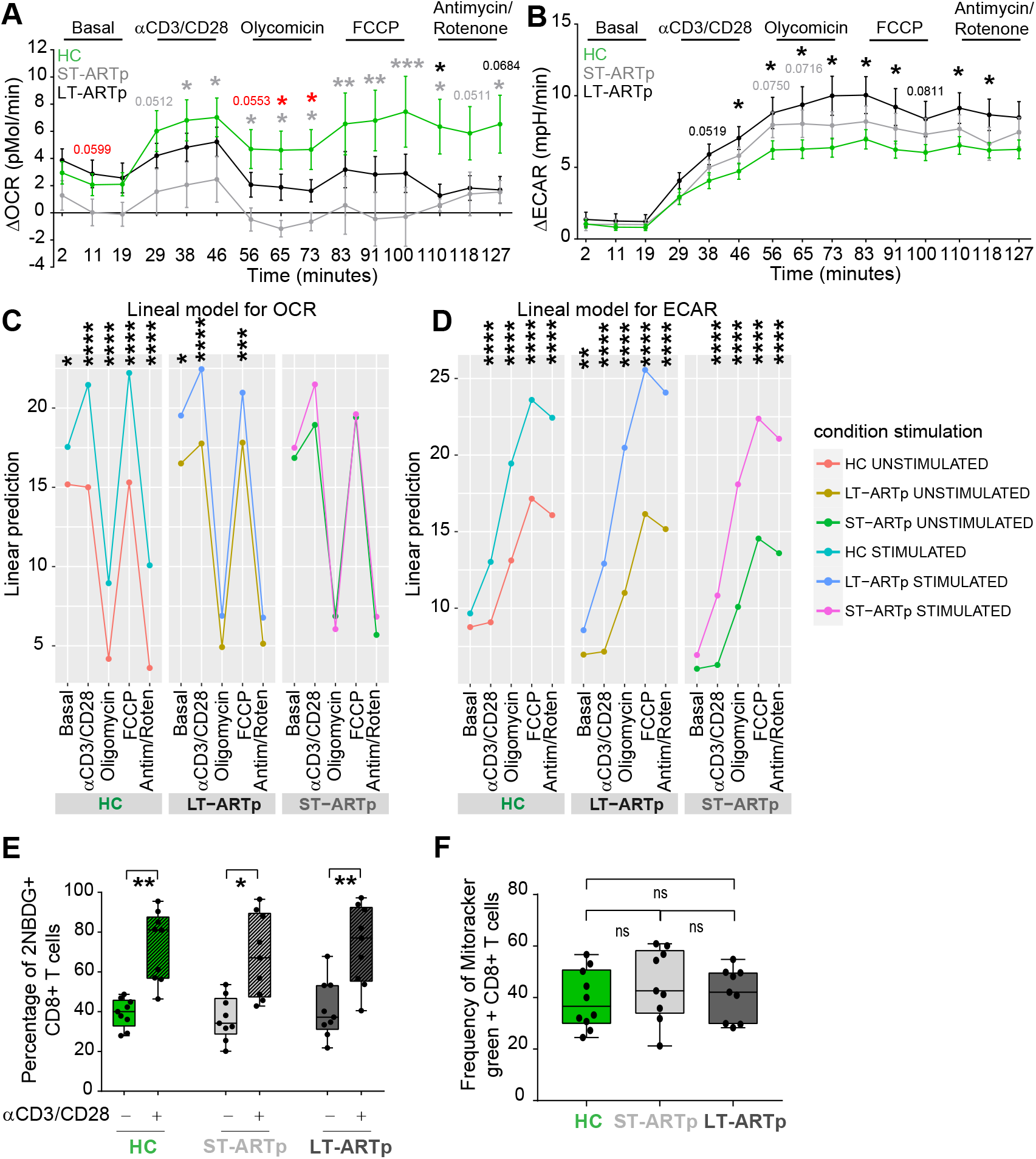
Metabolic analyses of CD8+ T cell from HC, ST-ARTp and LT-ARTp. (A-B): Delta of OCR (A) and ECAR (B) between TCR activated *vs* basal levels on CD8+ T cells from HIV negative (HC), ST-ARTp and LT-ARTp. Statistical significance per each timepoint was calculated with a multiple t test analysis. Light gray values show significance between ST-ARTp and HC; dark gray values show significance between LT-ARTp and HC; and red values show significance between ST-ARTp and LT-ARTp. (C-D): Linear mixed model of analysis for OCR (C) and ECAR (D) showing prediction mean for each timepoint set of treatment comparing basal and activated CD8+ T cells from HC (left), LT-ARTp (middle) and ST-ARTp (right). Statistics of comparison were calculated with the package lmer from lmerTest on R version 4.1.1 (see methods). (E): Glucose uptake analyzed by percentage of cells incorporating fluorescent 2NBDG from isolated CD8+ T cells by flow cytometry comparing basal and TCR activated CD8+ T cells. Statistical significance was calculated using two-tailed Wilcoxon test (*p<0.05; **p<0.01). (F): Mitochondrial mass detection was determined by flow cytometry comparing MitoTracker straining on total CD8+ T cells from HC (green), ST-ARTp (light gray) and LT-ARTp (dark gray) used for metabolic analyses. Statistical significance between groups was calculated using two-tailed Mann-Whitney U test and statistical significance between memory subsets within the same group using two-tailed Wilcoxon test (*p<0.05; ****p<0.0001).

**Supplementary figure 8.**
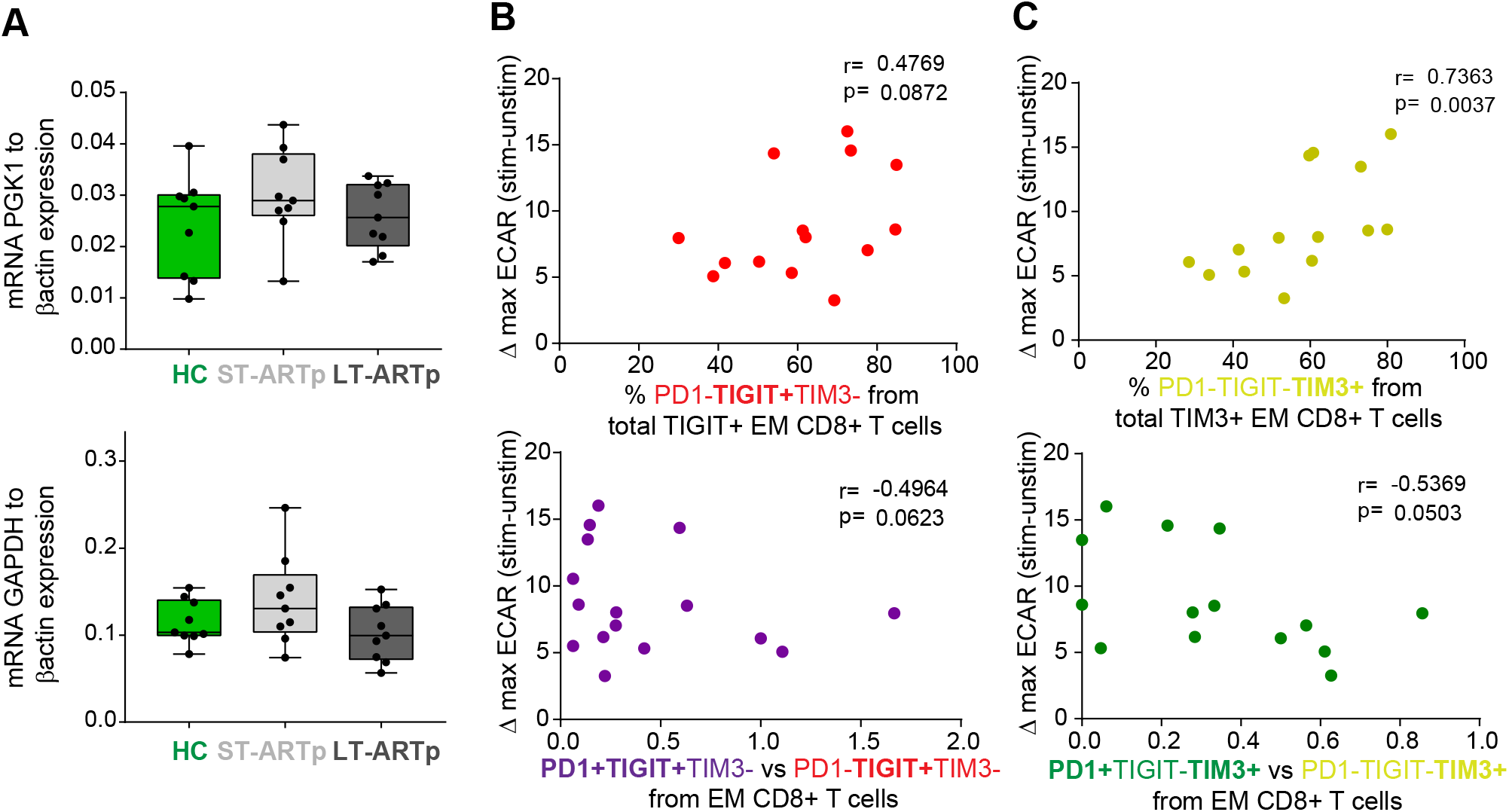
Transcription of different genes involved in glycolytic metabolism and correlation between ECAR and exhaustion patterns in memory CD8+ T cell from PLWH. (A): RT-qPCR analysis of PGK1 (upper) and GAPDH (lower) transcriptional expression normalized to β–actin mRNA levels in memory CD45RA-CD8+ T cells isolated from total PBMCs of HIV-1 negative controls (HC; green) and PLWH subgroups (ST-ARTp; light gray and LT-ARTp; dark gray). Statistical significance between double and single positive populations was calculated using two-tailed Wilcoxon test (*p<0.05). (B-C): Spearman correlations between ΔECAR at maximal activation after TCR stimulation (73 minutes) and proportions of PD1-TIGIT+TIM3-(B, upper panel; red) or PD1-TIGIT-TIM3+ (C, upper panel; yellow) to total TIGIT+ or TIM3+ cells or ratios of the indicated populations (lower plots; PD1 *vs* TIGIT purple and red; PD1 *vs* TIM3 green and yellow) within effector memory CD8+ T cells. Spearman r and p values are shown on the upper left area of each plot.

